# Oncogene Regulated Release of Extracellular Vesicles

**DOI:** 10.1101/2020.04.04.025726

**Authors:** Seda Kilinc, Rebekka Paisner, Roman Camarda, Olga Momcilovic, Rebecca A. Kohnz, Noelle D. L’Etoile, Rushika M. Perera, Daniel K. Nomura, Andrei Goga

**Affiliations:** Department of Cell & Tissue Biology, University of California, San Francisco, San Francisco, CA 94143, USA; Biomedical Sciences Graduate Program, University of California, San Francisco, San Francisco, CA 94143, USA; Departments of Chemistry and Nutritional Sciences and Toxicology, University of California, Berkeley, Berkeley, CA 94720, USA; Department of Anatomy and Helen Diller Family Comprehensive Cancer Center, University of California, San Francisco, San Francisco, CA 94143, USA; Department of Medicine, University of California, San Francisco, San Francisco, CA 94143, USA; Deparment of Biology, Johns Hopkins University, Baltimore, MD 21218, USA

**Keywords:** oncogenes, extracellular vesicles (EVs), ceramide, ESCRT, MYC, HRAS, AURKB, biomass, lysosome

## Abstract

Oncogenes can alter cellular structure, function, development and metabolism including changing the balance between anabolic and catabolic processes. However, how oncogenes regulate tumor cell biomass remains poorly understood. Using isogenic mammary breast epithelial cells transformed with a panel of ten oncogenes found commonly mutated, amplified or overexpressed in multiple cancers, we show that specific oncogenes reduce the biomass of cancer cells by promoting extracellular vesicle release. While MYC and AURKB elicited the highest number of EVs, each oncogene tested selectively altered the protein composition of released EVs. Likewise, miRNAs were differentially sorted into EVs in an oncogene-specific manner. MYC overexpressing cells require ceramide, while AURKB require ESCRT to release high levels of EVs. Finally, lysosome-associated genes are broadly downregulated in the context of MYC and AURKB, suggesting that cellular contents instead of being degraded, were released via EVs. Thus, oncogene mediated biomass regulation via differential EV release is a new metabolic phenotype which may have implications for cellular signaling and homeostasis.

## Introduction

Cancer cells reprogram metabolic processes to accommodate nutrient availability, energy needs and biosynthetic activity to support cell survival under stressful conditions or to increase biomass to support their proliferation (DeBerardinis, Chandel, 2016). Glycolysis and glutaminolysis are utilized by tumors to create metabolic intermediates that are used as precursors of macromolecule synthesis and energy production (DeBerardinis et al., 2008). Tumors frequently have mutations in PI3K and AKT oncogenes which result in aberrant activation of mTORC1 pathway that induce an anabolic growth program resulting in nucleotide, protein and lipid synthesis (Yuan, Cantley, 2008). Likewise, MYC increases anabolic growth by altering genes involved in glycolysis, fatty acid synthesis, glutaminolysis, and serine metabolism (Stine et al., 2015). When nutrients are limited, tumor cells activate catabolic pathways like fatty acid oxidation (FAO) to increase ATP levels (DeBerardinis, Chandel, 2016). MYC high triple negative breast cancer cells rely on FAO to fuel bioenergetic metabolism (Camarda et al., 2016). Likewise, intercellular proteins and other macromolecules can be recycled via autophagy to maintain pools of metabolic intermediates (Galluzzi et al., 2014). For example, in Kras- or Braf-driven non-small-cell lung cancer cells, the supply of glutamine, the fuel for mitochondria, is maintained through autophagy (Strohecker, White, 2014, Guo et al., 2011). Alternatively, cancer cells can internalize proteins and other macromolecules from the tumor microenvironment via macropinocytosis. For example, Kras-driven pancreatic cancer cells perform macropinocytosis to maintain their amino acid supplies (Commisso et al., 2013). In contrast, cells can lose their biomass through extracellular vesicle (EV) release. Biomass loss via EV release is a facet of cancer metabolism that has not been explored in the context of specific transforming oncogenes.

Though the link between cancer drivers and biomass regulation has not been well-explored, EVs have been characterized with respect to their size, biogenesis and content. Extracellular vesicles are secreted lipid bilayer membrane enclosed vesicles which contain proteins, lipids, RNA and DNA (Minciacchi, Freeman & Di Vizio, 2015). Every cell secretes heterogenous populations of EVs ranging in size and differing in their biogenesis. Largest in size are the >1000nm apoptotic bodies secreted from dying cells, which contain histones and fragmented DNA. The 100-1000nm, large EVs, generally termed microvesicles (MVs), are formed by blebbing of plasma membranes. Exosomes, small 30-150nm EVs originate by inward budding of endosomal membranes into multivesicular bodies (MVBs) (Bebelman et al., 2018, Gustafson, Veitch & Fish, 2017, Colombo, Raposo & Thery, 2014) and are released as MVBs fuse with the plasma membrane. Interest in EV biogenesis stems from the possibility that the diverse contents of EVs act as messages that a cancer cell passes to the cells in its environment (Desrochers, Antonyak & Cerione, 2016).

Several studies have indicated that cancer cells release EVs which do indeed reshape the tumor microenvironment by triggering angiogenesis, permitting immune surveillance escape or reprogramming behavior of surrounding cells (Poggio et al., 2019, Kalluri, 2016, Becker et al., 2016, Desrochers, Antonyak & Cerione, 2016). Recent years have seen an increase in the understanding of the various cargos and roles of EVs in cancer development and progression and their potential use as biomarkers or vehicles of drug therapy (EL Andaloussi et al., 2013, Sousa, Lima & Vasconcelos, 2015, Kalluri, 2016). miRNAs are frequently identified EV cargos which may function to reprogram the recipient cells, for example exosomal miRNAs secreted from salivary mesenchyme regulate epithelial progenitor expansion during organogenesis (Hayashi et al., 2017). Similarly, exosomal miRNAs secreted by metastatic breast cancer cells target tight junction protein and destroy vascular endothelial barriers of surrounding cells promoting migration and metastasis (Zhou et al., 2014). The crosstalk from stromal to breast cancer cells via exosomes has also been shown to regulate therapy resistant pathways (Nabet et al., 2017, Luga et al., 2012). Alternatively, cancer cells can utilize EVs to secrete tumor suppressive miRNAs and toxic lipids to favor tumor growth (Kanlikilicer et al., 2016). Though several oncogenes have been shown to play a role in altering the content of extracellular vesicles and the functional effects of these EVs on the recipient cells, how different oncogenes regulate EV biogenesis and release remains poorly understood.

EV production within MVBs can be promoted by at least two well-characterized pathways: one that is ceramide dependent and the other that uses the ESCRT machinery. Ceramide lipids change the curvature of the membrane and can be locally concentrated within endosomal membranes by the action of neutral sphingomyelinase. High concentrations of ceramides within a membrane, trigger their inward budding thus forming an EV-filled MVB (Trajkovic et al., 2008). Membrane budding and EV production within an MVB can also be triggered by the <u>E</u>ndosomal <u>S</u>orting <u>C</u>omplex <u>R</u>equired for <u>T</u>ransport (ESCRT) which is comprised of four subcomplexes; ESCRT-0 which recognizes and clusters ubiquitinated proteins; ESCRT-I and -II which deform endosomal membranes to form buds where cargo is sorted. Finally, ESCRT-III together with accessory proteins cleave the buds to form intraluminal vesicles (ILVs) (Wollert, Hurley, 2010, Henne, Buchkovich & Emr, 2011, Colombo, Raposo & Thery, 2014)The MVBs are then trafficked to the plasma membrane and once they fuse with the plasma membrane, their contents, the exosomes, are released to the surrounding environment (Colombo et al., 2013, Baietti et al., 2012). While oncogenes can reprogram various aspects of cellular processes like proliferation, metabolism, epithelial to mesenchymal transitions, little is known about how they affect extracellular vesicle biogenesis, heterogeneity and release. Using an isogenic panel of oncogene-transformed cells, we examine quantitative and qualitative effects of EV release induced by different oncogenes.

We find that distinct oncogenes regulate the biogenesis and release of different amounts and sizes of EVs. EVs released from transformed cells also demonstrate distinct protein and miRNA content dependent on the driver oncogenes. This implies that oncogenes regulate cellular biomass release through EVs a new metabolic phenotype of cancer cells.

## Methods

### Cell Lines

MCF10A isogenic cell lines were stably generated using retroviral infection in our previous study (Martins et al., 2015). Control MCF10A cell line was generated using empty vectors of puromycin and blasticidin (Martins et al., 2015). All MCF10A cell lines were grown according to published protocols (Debnath et al., 2002). The EC4 conditional liver tumor line used in Figure 2 was a gift of D. Felsher at Stanford University. EC4 cells were grown in high glucose DMEM supplemented with 10% FBS and 1X non-essential amino acids. We add 8ng/ml doxycycline to the media to turn off MYC expression (Anderton et al., 2017). RPE-NEO and RPE-MYC cells were a gift from J. Michael Bishop and cultured as described (Goga et al., 2007).

**Figure 1.**
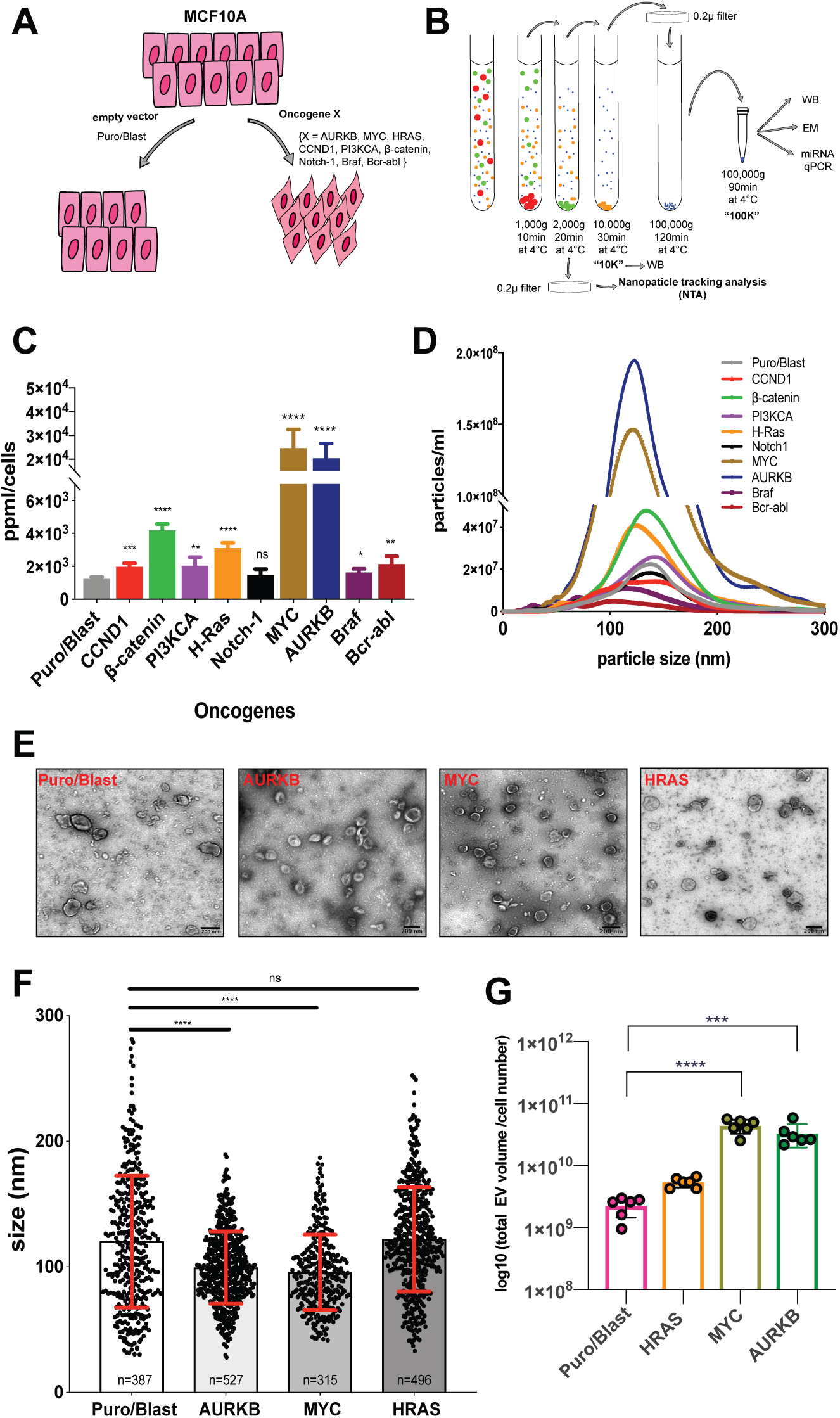
Oncogenes alter abundance and size of released EVs. **A.** Overview of MCF10A isogenic cell line system.10 different MCF10A cell lines generated by transducing with empty vectors as control (i.e.; Puro/Blast) or with a given oncogene X. **B.** Schematic representation of extracellular vesicle isolation protocol anddownstream analysis.**C.** Nanoparticle tracking analysis (NTA) of extracellular vesicles isolated from MCF10A oncogenic lines. Particle per ml (ppml) counts normalized to cell number (ppml/cells) for each oncogenic line. NTA is performed 3 times 30 second for each sample. n=6, ns p>0.05, * p<0.05, ** p< 0.01, *** p<0.001, **** p<0.0001.**D.** Size distribution of extracellular vesicles isolated from MCF10A oncogenic lines. Each oncogenic line analyzed with NTA (panel C) and mean values of 6 different analysis are plotted. **E.** Transmission electron micrograph (TEM) of purified EVs(100K fractions) of Puro/Blast, HRAS, AURKB and MYC overexpressing MCF10A lines. Scale bar 200nm. **F.** Size quantification of purified EVs from TEM images using Fiji. TEM repeated two times and representative random images from two different experiments were analyzed. Total number of vesicles analyzed are shown below each bar. ns p>0.05, ****p<0.0001. **G.**The biomass loss was calculated as total volume of released EVs normalized to cell number. Calculations were done using NTA data from panel C and D. n=6, *** p<0.001, **** p<0.0001.

**Figure 2.**
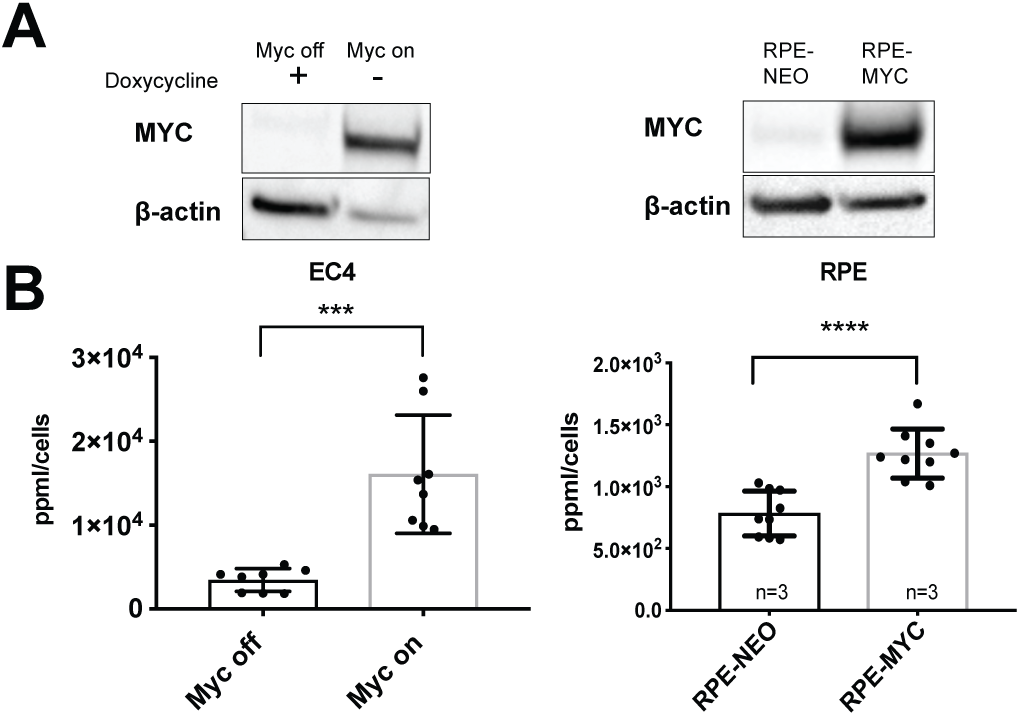
MYC overexpression also produces high number of EVs in other cell types. **A.**Western blot analysis of MYC overexpression in EC4 (left) and RPE (right) cells. **B.** Nanoparticle tracking analysis of EC4 (liver tumor) and RPE (retinal pigmented epithelial) cells. Particles per ml(ppml) are normalized to cell number (ppml/cells) and experiments repeated 3 times with two replicates for each line. ***p<0.001, ****p<0.0001.

### Nanoparticle Tracking Analysis (NTA)

We seeded 2×10^5^ cells / well into 6 well dishes and replaced media with 5% KSR (Knockout Serum Replacement) (Gibco) containing media the next day. We cultured cells in 5% KSR media for 48 hours. The conditioned media was collected and centrifuged at 1000g spin for 10 mins. Then supernatant was centrifuged at 2000g for 20mins. The supernatant from 2000g spin was filtered through a 0.2μ filter. We performed nanoparticle tracking analysis (NTA) using NanoSight LM10 (Malvern Panalytical). We kept all the settings (gain, camera level, and thresholds) for capturing the videos and analysis constant for all samples. We measured each sample 3 times for 30 seconds. We used Nanoparticle Tracking Analysis Software 3.2. For each sample, cells were collected, stained with Trypan Blue Stain (0.4%) (Invitrogen) and counted using Countess Automated Cell Counter (Invitrogen) according to the manufacturer’s instructions to determine viable cell number. We normalized each particle count to the cell number.

### EV isolation

The same number of cells were seeded, and next day media was replaced with 5% KSR after washing with twice with PBS. EVs were isolated using differential centrifugation after 48 hours of media change (Figure 1B). Briefly, the condition media collected and centrifuged at 1000g for 10 mins at 4°C to pellet cells. Supernatant was centrifuged at 2000g for 20min at 4°C, transferred to the new tubes and centrifuged at 10,000g for 30 min at 4°C (pellet of this spin was 10K fraction). Supernatant was transfer to a Beckman tube and centrifuged at 100,000g using SW28 rotor (Beckman). Supernatant was removed leaving 1ml behind. The remaining 1ml supernatant was then mixed with pellet and transferred to a Beckman Eppendorf tube and re-centrifuged at 100,000g using rotor TLA-100.3 (pellet of this spin is 100K fraction). Cells from the first 1000g pellet were pooled with cells trypsinized from the plates and counted by Countess (Invitrogen) using Trypan Blue stain 0.4% (Invitrogen). EVs were isolated from three 10cm plates for determining the protein composition of EVs (Figure 3). For GW4869 and siRNA knockdowns, one 10cm plate was used for EV isolation (Figure 5 and 6).

**Figure 3.**
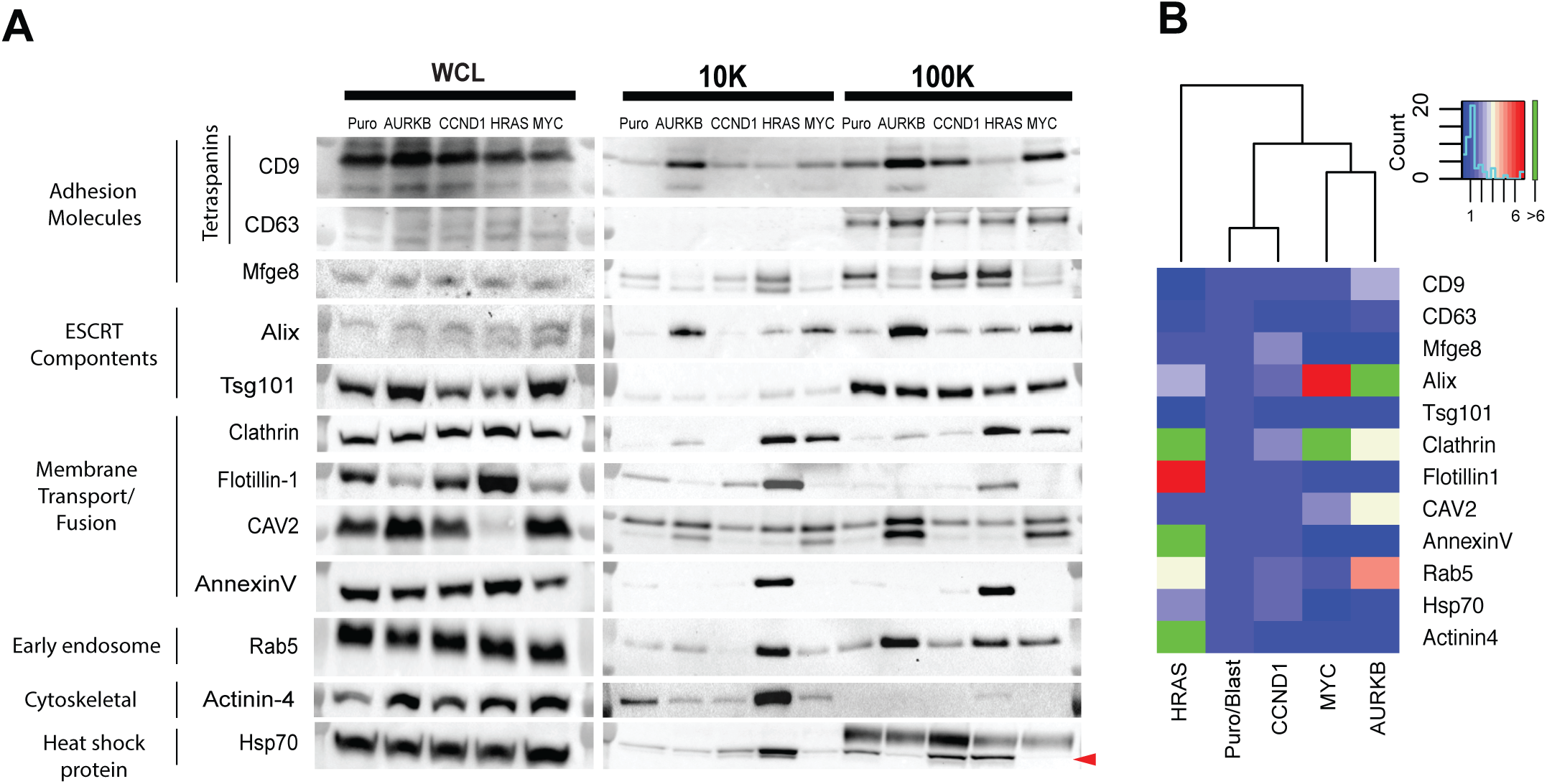
Oncogenes alter protein composition of extracellular vesicles. **A.**Western blot analysis of selected EV markers on whole cell lysates (WCL) on the left, 10K and 100K fractions of purified EVs on the right. EVs are isolated from control (Puro/Blast), AURKB, CCND1, HRAS and MYC overexpressing MCF10A cells. One representative image of three independent experiments is shown. Red arrow indicates expected band size. **B.** Quantification of three independent western blot analysis of panel A is illustrated as heat map. Protein levels in each oncogenic EVs are normalized to the levels of control (Puro/Blast) EVs. Average values of three independent experiments are depicted in the heat map.

**Figure 4.**
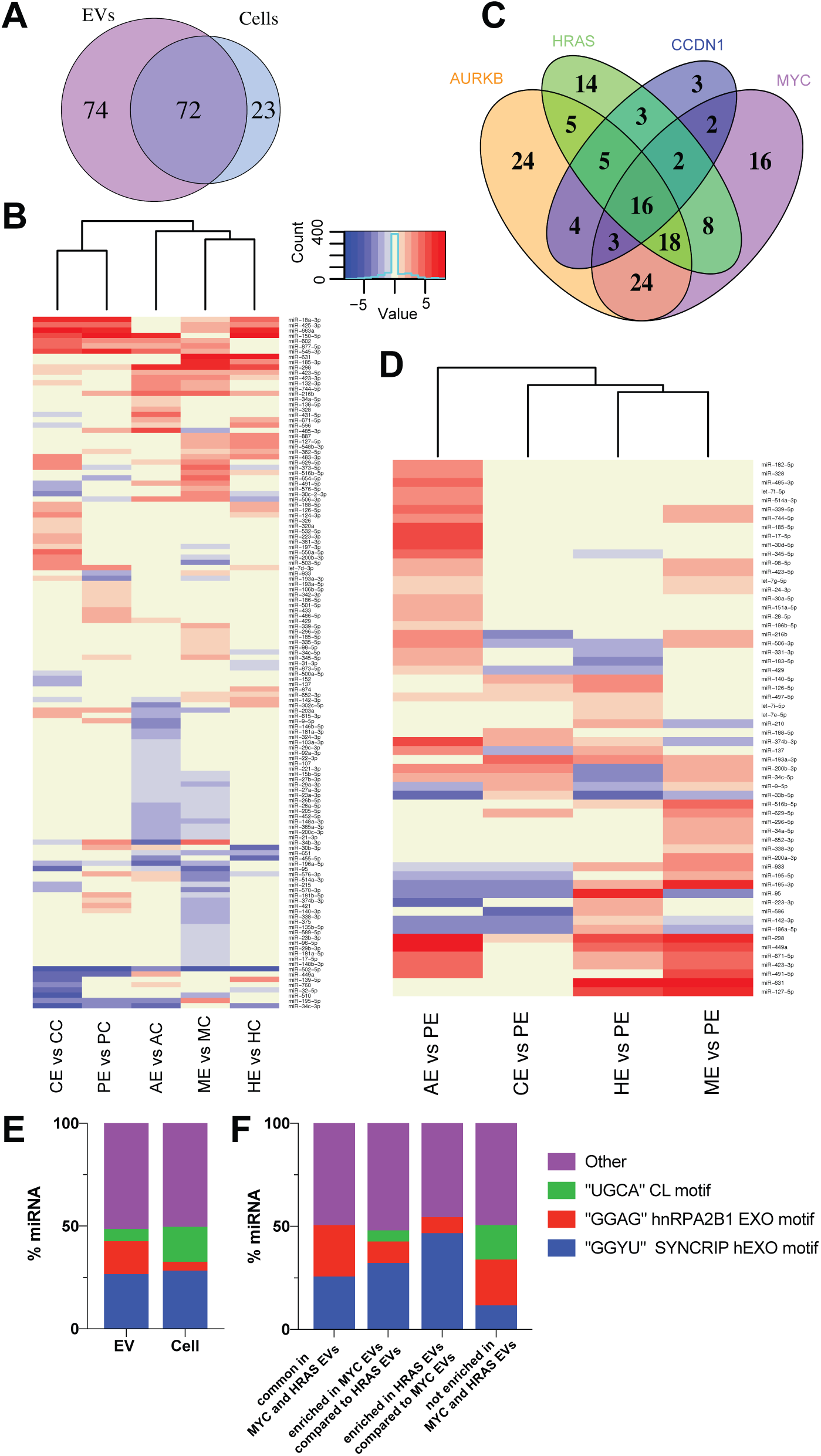
Oncogenes differentially sort miRNAs into EVs. **A**. Venn diagram of differentially regulated miRNAs in EVs and cells from AURKB, CCND1, HRAS and MYC overexpressing MCF10As. **B.**Heatmap analysis of altered miRNAs in EVs versus corresponding cells. Each column represents a given oncogenic line and rows represent identified miRNAs. Downregulated miRNAs in EVs represented in blue, whereas upregulated miRNAs in EVs depicted in red. CE=CylinD1 EV, CC=CylinD1 cells, PE=Puro/Blast EVs, PC=Puro/Blast cells, AE=AURKB EVs, AC=AURKB cells, HE=HRAS EVs, HC=HRAS cells, ME= MYC EVs, MC=MYC cells.**C.** Venn diagram of shared and distinct miRNAs in oncogenic EVs that are dysregulated compared to control EVs. **D.** The upregulated miRNAs compared to control EVs in at least one of the oncogenic EVs are presented as heat map. **E.**Histogram showing the percentage of miRNAs with known motifs in cell vs EVs. **F.** Histogram showing the percentage of miRNAs with known motifs in HRAS and MYC EV groups.

**Figure 5.**
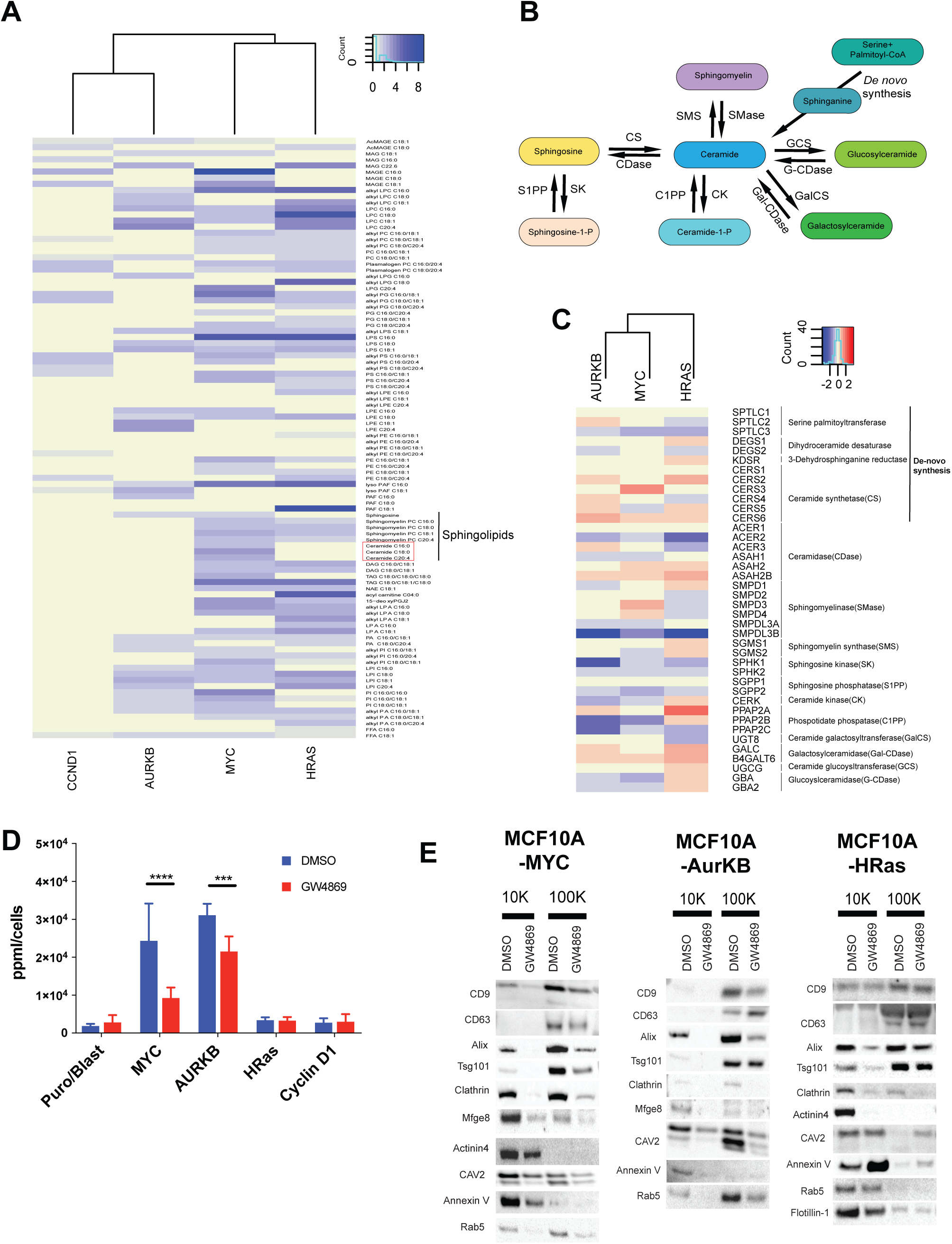
MYC overexpression alters ceramide metabolism and results in high number of EV release. **A.**Fold change in lipid levels from oncogene-expressing MCF10A cells vs control cells is represented as heat map. Only significantly (p < 0.05 using student t-test) dysregulated lipids are depicted in the heat map. 6 different samples are used in each oncogene-expressing group. **B.** Schematic representation of ceramide biogenesis.**C.** Heat map analysis of altered genes in the ceramide metabolism from RNA sequencing of AURKB, HRAS, MYC overexpressing MCF10A lines. **D.**Nanoparticle tracking analysis (NTA) of EVs from MCF10A cells either treated with vehicle (DMSO) or neutral sphingomyelinase inhibitor, GW4869 (5μM) for 48 hours. Each MCF10A line is repeated at least 5 times. Particles per ml are normalized to cell number (ppml/cells) in each group. NTA is performed 3 times 30 second for each sample. ***p<0.001, ****p<0.0001.**E.** Representative western blots of EV markers on 10K and 100K EV fractions from MCF10A cells that are either treated with vehicle (DMSO) or 5μM GW4869.

**Figure 6.**
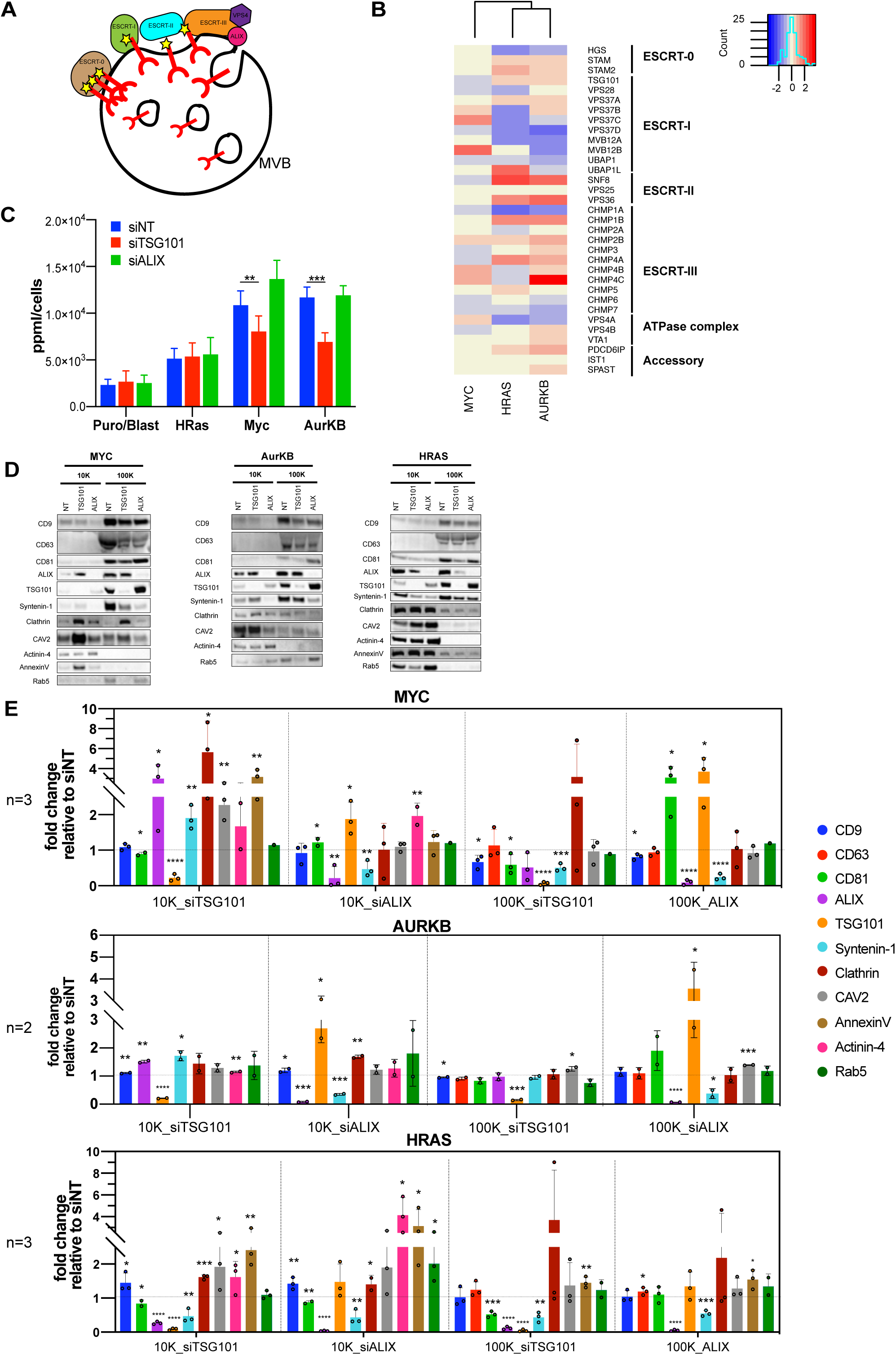
TSG101 is required for increased EV production in AURKB and MYC cells. **A.** Schematic representation of ESCRT-dependent ILV budding in MVBs. **B.**Heat map representation of dysregulated genes in ESRCT pathway from RNA sequencing of AURKB, MYC and HRAS overexpressing cells. **C.** Nanoparticle tracking analysis (NTA) of EVs from MCF10A cells that are transfected with siRNA pools of non-targeting (siNT), TSG101 (siTSG101), and Alix (siALIX). Each experiment repeated three times with two replicates. Particle per ml counts were normalized to cell number (ppml/cells) in each group. NTA is performed 3 times 30 second for each sample. **p< 0.01, ***p<0.001. **D.** Representative western blots of EV markers on 10K and 100K EV fractions from MCF10A cells that are transfected with siRNA pools of non-targeting (siNT), TSG101 (siTSG101), and Alix (siALIX). **E.** Quantification of western blot analysis from D. Three independent experiments performed for MYC and HRAS, two for AURKB. Quantification of protein bands is performed using ImageLab software from BioRAD. Fold change of EV markers relative to siNT treatment in each group compared and each value depicted in the bar graph.

### Electron Microscopy

∼5-6μl of the resuspended 100,000 ×g pellet fraction was spread onto glow discharged Formvar-coated copper mesh grids (Electron Microscopy Sciences) and stained with 2% Uranyl acetate for 2 min. Excess staining solution was blotted off with filter paper. After drying, grids were imaged at at 120 kV using a Tecnai 12 Transmission Electron Microscope (FEI, Hillsboro, OR). EV size analysis on micrographs were done using Fiji software.

### EV and cell volume calculations

We assumed released EVs as spherical particles and performed a biomass calculation as follows: 1) we calculated the volume of the sphere (vesicle) by each size range, 2) we multiplied the frequency (number) of the vesicles by each size range, 3) we summed all size ranges to get a total volume of the released biomass, 4) we divided total volume by cell number for normalization. We measured cell size using Countess automated cell counter (Invitrogen). We measured each cell line at least four times and take average of measured cell sizes. We assumed cells as spheres and calculated their volume.

### Western Blot Analysis

Cells, 10K and 100K pellets were lysed in Pierce RIPA buffer (25mM Tris-HCl (pH 7.6), 150mM NaCl, 1% NP-40, 1% sodium deoxycholate, 0.1% SDS) supplemented with protease and phosphatase inhibitor cocktail (Roche) on ice for 20 mins. Cell pellets were spun down at 13,000g for 15 mins at 4°C. Protein concentration was determined using the DC Protein Assay (BioRad). For whole cell lysates, 30 μg protein extracts were resolved using 4–12% SDS-PAGE gels (Life Technologies) and transferred to nitrocellulose membranes using iBlot (Life Technologies). For 10K and 100K fractions, the same number of cells were seeded at the beginning of each experiment, and EV pellets were resuspended in the same volume of RIPA buffer and same volume loaded for SDS-PAGE. Membranes were probed with primary antibodies overnight on a 4°C shaker, then incubated with horseradish peroxidase (HRP)-conjugated secondary antibodies, and signals were visualized with Visualizer™ Western Blot Detection Kit (Millipore). Band analysis was performed using ImageLab software from BioRAD and normalized to cell counts.

### RNA isolation and qPCR

RNA from cells and EVs was isolated using mirRNeasy micro kit (Qiagen). cDNA was prepared using Universal cDNA synthesis kit according to manufacturer’s instructions (Exiqon/Qiagen). qPCR was performed using miRCURY LNA miRNA Human Panel I according to manufacturer’s instructions (Exiqon/Qiagen). The data was analyzed using GenEX software (MultiD) according to manufacturer’s instructions. Normalization was performed using NormFinder tool of the software. Three independent replicates were used for MYC and Puro/Blast cell and EV samples. Two independent replicates were processed for HRAS, CCND1, AURKB cell and EV samples.

### RNA sequencing

RNA was isolated using RNeasy mini kit according to manufacturer’s instructions (Qiagen). Three different samples of RNA were isolated for each MCF10A line (i.e.; Puro/Blast, MYC, AURKB and HRAS). Library preparation and Illumina sequencing were performed at Novogene Corporation (Sacramento, CA) (en.novogene.org). Briefly, mRNA from Eukaryote organisms was purified from total RNA using poly-T oligo-attached magnetic beads. The mRNA was first fragmented randomly by addition of fragmentation buffer, then NEB library was prepared. Q-PCR is used to accurately quantify the library effective concentration (> 2nM), in order to ensure the library quality. Libraries fed into Illumina Platform, and paired-end reads were generated. Downstream analysis was performed using a combination of programs including STAR, HTseq, Cufflink and Novogene’s wrapped scripts. Alignments were parsed using Tophat program and differential expressions were determined through DESeq2/edgeR. Heat maps were generated with the gplots package in R (version 3.3.1).

### Metabolomic profiling

Metabolomic analyses were conducted as previously described in Louie et. al. (Louie et al., 2016). Briefly, 2 million cells were plated overnight, serum starved for 2 hours prior to harvesting, after which cells were washed twice with PBS, harvested by scraping, and flash frozen. For nonpolar metabolomic analyses, flash frozen cell pellets were extracted in 4mL of 2:1:1 chloroform/methanol/PBS with internal standards dodecylglycerol (10 nmoles) and pentadecanoic acid (10 nmoles). Organic and aqueous layers were separated by centrifugation, and organic layer was extracted. Aqueous layer was acidified with 0.1% formic acid followed by re-extraction with 2 mL chloroform. The second organic layer was combined with the first extract and dried under nitrogen, after which lipids were resuspended in chloroform (120 μl). A 10 μl aliquot was then analyzed by both single-reaction monitoring (SRM)-based LC-MS/MS or untargeted LC-MS. For polar metabolomic analyses, frozen cell pellets were extracted in 180 μl of 40:40:20 acetonitrile/methanol/water with internal standard d3 N15-serine (1 nmole). Following vortexing and bath sonication, the polar metabolite fraction (supernatant) was isolated by centrifugation. A 20 μl aliquot was then analyzed by both single-reaction monitoring (SRM)-based LC-MS/MS or untargeted LC-MS. Relative levels of metabolites were quantified by integrating the area under the curve for each metabolite, normalizing to internal standard values, and then normalizing to the average values of the control groups.

### GW4869 treatment

1×10^5^ cells seeded into one well of 6well plates at the beginning of experiment. The media changed to 5% KSR with 5μM GW4869 (Sigma) or vehicle, DMSO (Sigma) on the next day. Condition media collected after 48 hours and NTA was performed as described above. For western blot experiments, 7.5×10^5^ cells were seeded into one 10cm dish.

### RNAi knockdown

*TSG101*-specific (L-003549-00-0005), *ALIX*-specific (L-004233-00-0005) and nontargeting (D-001810-10-20) siRNAs were purchased from GE Dharmacon (SMARTpool, four siRNAs per gene). 30 pmol siRNA was used to transfect cells with the Lipofectamine RNAiMAX Transfection Reagent (Life Technologies), according to the manufacturer’s instructions. Cells were incubated with siRNA and at 24 hrs media was changed to 5%KSR. Cells were incubated for additional 48hrs to collect EVs. At the beginning of experiments, 1×10^5^ cells seeded per well of 6-well plates for NTA analysis and 7.5×10^5^ cells per 10cm dish for western blot analysis. At the end point, cells were collected, stained with Trypan Blue Stain (0.4%) (Invitrogen) and counted using Countess Automated Cell Counter (Invitrogen) according to the manufacturer’s instructions. Particle counts and band intensities are normalized to the cell number.

## Results

### Oncogenes alter EV abundance and size

We sought to understand how different oncogenes alter the number and contents of EVs released from transformed cells. MCF10A cells are derived from non-tumorigenic breast tissue but are susceptible to transformation by a wide range of oncogenes, allowing us to study diverse oncogene signaling pathways (Debnath et al., 2002, Martins et al., 2015). We generated a panel of isogenic MCF10A human mammary epithelial cell lines engineered to express a variety of the most commonly mutated, amplified or overexpressed oncogenes in breast and other types of cancers (Martins et al., 2015). In order to directly compare the influence of specific oncogenic signals on EV properties we used a panel of 10 cell lines in which individual oncogenes were stably expressed (Martins et al., 2015) and compared these to cells transduced with empty vectors (i.e., Puro/Blast) (**Figure 1A**).

EVs were isolated using a differential ultracentrifugation method as depicted in **Figure 1B** (Muralidharan-Chari et al., 2009, Thery et al., 2006). Prior to EV collection, each cell line was cultured for 48 hours in defined medium (KSR) that does not contain exogenous EVs, normally abundant in serum-containing media. This allowed us to ensure that EVs were derived from each of the isogenic MCF10A lines. To assess the number and size of EVs produced we utilized two complementary assays. First, we performed nanoparticle tracking analysis (NTA) on 10 different isogenic MCF10A lines using NanoSight video microscopy tracking (**Figure 1C-D**). We found that 8 out of 9 oncogene-expressing cell lines significantly increased the number of particles released compared to control (Puro/Blast) cells (**Figure 1C**). Amongst the oncogenes tested, MYC and AURKB-overexpressing MCF10A cells release many more EVs (∼ 20.5- and 16.9-fold respectively) than control cells, and also significantly higher than any of the other oncogene-expressing cell lines (**Figure 1C**). Furthermore, the size distribution of the EVs revealed that the majority of released vesicles are small EVs (sEV) <200nm in size (**Figure 1D**) consistent with known size distribution for exosomes (Colombo, Raposo & Thery, 2014). MYC and AURKB-derived EVs were smaller with a modal particle size of 116.9 nm and 119.1 nm, respectively **(Supplementary Figure 1).** Thus, we observed that most oncogenes induced significantly increased EV release, though MYC and AURKB vesicles were far more numerous on a per cell basis but tended to be smaller (**Figure 1C-D**).

NTA analysis does not permit direct visualization of individual vesicle morphology. We therefore used a second approach, electron microscopy (EM), to address qualitative differences and confirm size changes via direct visualization of small EVs. We used transmission electron microscopy to image small EVs collected from the 100K fraction (**Figure 1B**) from MYC, AURKB and HRAS(G12V) oncogene expressing cells and control MCF10A cells. We observed the typical cup-shaped morphology previously described for small EVs **(Figure 1E)** (Sahoo et al., 2011). To elucidate the size differences of EVs we measured the diameter of each EV by random sampling from micrographs. We analyzed a minimum of 300 EVs per condition. The mean size of control (Puro/Blast) EVs is 120nm whereas the mean size of EVs from AURKB and MYC overexpressing cells are 99nm and 96nm respectively. HRAS EVs were of similar size and appearance to Puro/Blast EVs with a mean size of 122 nm. MYC and AURKB EVs showed smaller size variance (sd ± 29, ±30nm, respectively) compared to control and HRAS EVs which demonstrated a wider size distribution (sd ± 52, ±42nm, respectively) **(Figure 1F).** These data indicate that different oncogenes alter the size but not morphology of released EVs and that MYC and AURKB oncogenes decrease the size of released EVs.

Since oncogenes induced differences in both the size and number of secreted EVs, we asked if the total biomass of released EVs was altered across the panel of oncogenes. Using the number of EVs released and their size distribution from NTA analysis, we determined the volume of cellular biomass released as EVs and normalized to total cell number. We found that MYC and AURKB expressing cells released ∼19.5- and ∼ 14.6-fold more biomass compared to control MCF10A cells respectively **(Figure 1G)**. We next measured cell volume to estimate percentage of cellular biomass released by EVs during a 48hr period of culture. We measured cell sizes using Countess automated cell counter and calculated cell volume using a spherical model. Control and HRAS overexpressing MCF10A cells released less than 1 % of their cellular volume whereas MYC and AURKB expressing MCF10As released 7.7 % and 4.1% of their total cellular volume via EVs (**Supplementary Figure 1B for cell sizes**). Remarkably, the volume of cells expressing either oncogene is within the same order of magnitude as control cells. Thus, MYC and AURKB overexpressing cells can significantly alter their biomass flux via EV release compared to control MCF10A cells to maintain cellular size homeostasis.

### MYC overexpression alters EV production in other cell types

MCF10A cells which overexpress MYC demonstrated the greatest number of EVs and EV biomass released. MYC is a pleiotropic transcription factor that alters transcription of many genes involved in a wide variety of processes, including but not limited to, proliferation, apoptosis and metabolism (Meyer, Penn, 2008, Camarda, Williams & Goga, 2017, Gabay, Li & Felsher, 2014). The MYC oncogene is also overexpressed in many of the most aggressive types of human cancers, such as lymphoma, receptor triple negative breast cancer, and liver cancer (Lin et al., 2012, Horiuchi, Anderton & Goga, 2014). We thus sought to determine if MYC overexpression affects EV production of other cell types. First, we asked if high EV production requires MYC over expression. We used EC4 cells derived from a conditional MYC transgenic mouse model of liver cancer in which MYC is induced to drive oncogenesis (Anderton et al., 2017, Cao et al., 2011). When cells are grown in the presence of doxycycline, MYC transgene expression is rapidly inhibited (**Figure 2A**). When MYC expression was turned off for 2 days, EV production decreased 5-fold (**Figure 2B**). Thus, acutely inhibiting MYC expression can rapidly reprogram cells to decrease EV release. Second, we utilized human retinal pigment epithelial (RPE) cells that constitutively overexpress human MYC (RPE-MYC) or a control plasmid (RPE-NEO) (Goga et al., 2007). MYC overexpression significantly increased EV release from RPE cells as well (∼ 2-fold) **(Figure 2B)**. Thus, across multiple cell lines, MYC overexpression induces a cellular program that increases EV release.

### The protein content of EVs is regulated by the driver oncogene

We postulated that specific oncogenes may alter not only the size and number of EVs released, but also their contents. To elucidate the heterogeneity of EVs released from different oncogenes, we performed western blot analysis of known EV markers from five different MCF10A cell lines including control, AURKB, CCND1, HRAS and MYC overexpressing cells. We performed three independent EV isolations, (representative western blots are shown (**Figure 3A**); the results of three different western blot analysis is shown as heat maps (**Figure 3B**) (also shown as histograms (**Supplementary Figure 2**)). To compare protein composition from different vesicle sizes, we performed analysis both from 10K fractions (10,000g spin, heavier fraction, enriched in microvesicles) and 100K fractions (100,000g spin, lighter fraction, enriched in sEVs). We performed western blots for candidate proteins that represent several classes of EV-associated molecules.

Tetraspanins (i.e., CD9 and CD63) are enriched in sEVs, but can be found in other types of EVs as well (Crescitelli et al., 2013, Kowal et al., 2016). We found that CD9 was present in both the 10K and 100K fractions, whereas CD63 was only detected in the 100K fraction **(Figure 3A)**. AURKB-derived EVs were significantly enriched in CD9 tetraspanins **(Figure 3B)**. *MFGE-8*, another adhesion molecule found in microvesicles and exosomes (Bobrie et al., 2012), was significantly enriched in CCND1-derived EVs and observed in both fractions. We also examined ESCRT components, *TSG101* and *ALIX*, which are known to induce exosome biogenesis (Colombo et al., 2013, Baietti et al., 2012). *TSG101* protein abundance was significantly decreased in AURKB, HRAS, and MYC-derived EVs (**Supplementary Figure 2**) however, *ALIX* abundance was only enriched in MYC and AURKB EVs **(Figure 3A-B)**.

Clathrin dependent endocytosis is one of the major sources of early endosomes (Le Roy, Wrana, 2005, Raiborg, Rusten & Stenmark, 2003). MYC and HRAS EVs contain significantly increased levels of clathrin **(Figure 3)**. *CAV2* is a major component of caveolae and functions to stimulate endocytosis and membrane lipid composition (Le Roy, Wrana, 2005, Cheng, Nichols, 2016). MYC and AURKB cells express high levels of *CAV2* and their EVs are also highly enriched in *CAV2* **(Figure 3)**, however other oncogene expressing cells release similar levels of *CAV2* containing EVs compared to controls. Flotillin-1 is also localized to caveolae, is involved in vesicle trafficking, and has been reported to be present in EVs (Baietti et al., 2012, Kowal et al., 2016, Colombo, Raposo & Thery, 2014, Le Roy, Wrana, 2005). We find that flotillin-1 is highly expressed in HRAS cells and also within the 10K fraction (**Figure 3A**). Its presence in control and CCND1 10K fractions but was barely detectable in 100K fractions also strengthens the observation that it is not a small EV specific marker (Kowal et al., 2016). Annexin-V, which binds to phosphatidyl serine (PS) enriched membranes, is significantly enriched only in HRAS-derived EVs (**Figure 3A**).

HRAS- and AURKB-derived EVs contains high levels of an early endosome marker, *RAB5*, though it is enriched in different fractions (**Figure 3**). *RAB5* is enriched in 10K fractions in HRAS EVs, whereas it is released abundantly in 100K fractions from AURKB overexpressing cells (**Figure 3A, 5E, 6D**). Actinin-4 is an actin-binding cytoskeleton protein and identified as a large EV (10K) marker in recent studies (Kowal et al., 2016, Jeppesen et al., 2019). Actinin-4 is highly abundant in HRAS-derived EVs. Finally, heat shock protein *HSP70*, is enriched in CCND1- and HRAS-derived EVs.

These results demonstrate that each oncogene alters EV protein content differently. MYC and AURKB oncogenes result in the release of small EVs that are enriched in endosomal origin associated proteins (i.e. high levels of *ALIX* and *CD9*). However, HRAS cells release very distinct types of vesicles that are enriched in large EV markers like actinin-4, flotillin-1, but are also highly enriched for Annexin-V, Clathrin, and *RAB5*. Thus, while AURKB and MYC EVs have similar size and protein content, CCND1 EVs are more similar to control EVs in protein content. Finally, while HRAS EVs are similar in size to control MCF10A cells, their protein content significantly differs from the rest of the control and oncogene expressing cells examined **(Figure 4B)**. Thus, EV protein content differs between cells transformed by distinct oncogenic drivers; the EV protein content differences may be useful to distinguish the oncogenic driver associated with their biogenesis.

### Sorting of miRNAs into small EVs is differentially regulated by oncogenes

We next sought to determine whether the miRNA content of EVs depends on the oncogenic driver. We isolated equal amounts of RNA from EVs or the corresponding whole cells that released them. We then performed qPCR analysis using a panel of 384 miRNA primers. We identified 95 miRNAs (about 1/4 of the panel) that were differentially expressed in oncogene-expressing cells compared to control cells (**Figure 4A, Supplementary Table 1**). 72 of these miRNAs were also enriched in oncogene-derived EVs compared to control EVs, however there were an additional 74 miRNAs that were differentially sorted into oncogene-derived EVs despite not being dysregulated in cells (**Figure 4A, Supplementary Table 2**). We proceeded to perform two types of analyses on these data: 1. We asked if each oncogene caused either preferential sorting into, or exclusion from, an EV by examining the concentration of each miRNA in EVs and comparing that to the concentration of miRNAs within the cell of origin. 2. We asked if the microRNAs within EVs from each cell type differed from that of the control cells or each other. Our goals were to discover what microRNAs the cells may be preferentially releasing, possibly in order to maintain homeostasis or to influence cells in their environment. We also sought to understand what microRNA handling and sorting processes may be employed when each oncogene is driving cellular phenotypes.

First, we determined which miRNAs are enriched in EVs versus their cell of origin (**Figure 4B**). The number and type of miRNA that is differentially released into EVs is unique to each oncogene. Although the released versus cellular miRNA profiles look different for each oncogene, CCND1-overexpressing cells seem most similar to control cells in the hierarchical clustering (**Figure 4B**). Certain miRNAs (e.g.; hsa-miR-18-3p, hsa-miR-425-3p, hsa-miR-663a, has-miR-150-5p, has-miR-298) were not only enriched in several oncogene-derived EVs but also in control EVs suggesting their release is independent of oncogenic driver. On the other hand, there are certain miRNAs that are preferentially released into EVs in an oncogene-specific manner (**Figure 4B**).

Second, we compared miRNAs differentially regulated from oncogene-derived EVs to control EVs (**Figure 4C**). There were 16 deregulated miRNAs common to all oncogenes either released or retained compared to control EVs (**Figure 4C**). AURKB and MYC derived EVs shared the highest number of deregulated miRNAs, released or retained, suggesting a common propensity to release these miRNAs. This might indicate a common mechanism is used in each cell type.

Third, we identified miRNAs that are uniquely over or underrepresented in EVs from each oncogene-expressing cell line. CCND1 derived EVs have the lowest number of differentially sorted miRNAs (3), whereas EVs from AURKB, HRAS and MYC oncogene-expressed (i.e.; 24, 14, 16 miRNAs respectively) **(Figure 4C)**.

The EV-enriched miRNAs from this dataset can potentially be explored as oncogene specific EV markers. We selected miRNAs that are enriched in at least one of the oncogene-derived EVs and presented them in **Figure 4D** to visualize unique or shared miRNAs **(Figure 4D)**. miRNA-298 was the only miRNA that was overrepresented in all of the oncogene-derived EVs. There were several miRNAs that are unique to MYC, AURKB and HRAS EVs **(Figure 4D)**. For example, miR-95 is significantly enriched in HRAS EVs and miR-34a-5p and miR-195-5p, which are well-known tumor suppressors, is highly specific to MYC EVs. Our data strongly suggests that oncogenes bias which miRNAs are loaded into EVs. This could result from oncogene specific expression of the secretion machinery, localization of the microRNA handling apparatus or microRNA binding proteins.

Thus, we asked whether the RNA motifs found within the differentially sorted microRNAs might help us identify oncogene-specific microRNA binding proteins. We examined known miRNA motifs that have been shown to control sorting of miRNAs into EVs and asked if these motifs are enriched in miRNAs that are sorted into EVs in an oncogene specific manner. Recently, a common motif (hEXO “GGYU”) was found to be enriched in miRNAs sorted into hepatocyte EVs by the RNA binding protein *SYNCRIP* (*HNRPQ*) (Santangelo et al., 2016). Likewise, two different miRNA motifs were reported to influence sorting of miRNAs into T cell-derived EVs: the EXO motif “GGAG” has been shown to bind hnRNPA2B1 which is proposed to sort the microRNAs bearing this motif into EVs (Villarroya-Beltri et al., 2013); the motif (CL motif “UGCA”) which is abundant in miRNAs *retained* in cells, was also identified in the same study (Villarroya-Beltri et al., 2013).

We first asked if any of these motifs were found in miRNAs either preferentially released into EVs or preferentially retained in cells that were in common between all oncogene expressing cell lines (**Figure 4E**). These motifs were only detected in half of these miRNAs (**Figure 4E**). The hEXO SYNCRIP binding motif was equally represented (25%) in both miRNAs enriched in EVs and retained in cells (**Figure 4E**). Interestingly, hnRNPA2B1 EXO motif was observed in 16% of the miRNAs sorted into EVs whereas it was detected in only 4.3% of miRNAs retained in cells. In contrast, the CL motif was abundant in miRNAs retained in cells (17%) while only 6% of the miRNAs enriched in EVs contain the CL motif (**Figure 4E, Supplementary table 3**). Although the SYNCRIP motif was detected in more miRNAs, hnRNPA2B1 motif seems to be more EV specific in MCF10A oncogene expressing cells. It is also noteworthy that half of the miRNAs either differentially loaded into EVs or retained in cells do not contain any of the previously reported sorting motifs. This suggests that miRNA EV sorting or cellular retention may occur via RNA binding proteins and/or mechanisms not previously described.

To further investigate oncogene-specific sorting, we analyzed miRNAs from MYC and HRAS derived EVs and examined the prevalence of previously identified sorting motifs. We categorized miRNAs that are common to both oncogenes, enriched in one of them compared to other, and not enriched in either (**Figure 4F**). miRNAs that are enriched in EVs common to both MYC and HRAS had equal representation (∼25%) for both the SYNRICP and hnRNPA2B1 motifs (**Figure 4F**). However, the SYNCRIP motif was more abundant (46%) in miRNAs that are enriched in HRAS EVs compared to MYC EVs (**Figure 4F, Supplementary table 3**). Interestingly, miRNAs not enriched in HRAS or MYC EVs (i.e.; enriched in AURKB and/or CCDN1 EVs) are two times more likely to have the hnRNPA2B1 motif (22%) compared to SYNCRIP motif (11%). Hence, miRNAs sorted into HRAS derived EVs appear to contain the SYNCRIP motif more frequently than other oncogenes tested. In aggregate, the microRNA sorting machinery may be differentially employed by each cancer driver.

We also sought to determine if miRNAs enriched in EVs are associated with oncogenic or tumor suppressive functions (**Supplementary Table 5**). For example, of the 6 miRNAs that were preferentially secreted via EVs from MYC over expressing cells as compared to cells expressing the other oncogenes, all six have been categorized as targeting anti-oncogenes or pro-growth processes, thus functioning as tumor suppressive miRNAs ((Wang et al., 2010); **Supplemental Table 5**). This may indicate that MYC high cells eliminate growth-suppressive miRNAs via preferential EV release. In contrast, many of the HRAS cells preferentially release miRNAs that are pro-tumorigenic (**Supplemental Table 5**), perhaps stimulating the growth of cells within the tumor microenvironment.

### Sphingolipid metabolism is altered for multiple oncogene transformed cells, but especially in MYC overexpressing cells

Exosomes originate from inward budding of early endosomal membranes sequestering lipids, proteins and RNA into intraluminal vesicles (Colombo, Raposo & Thery, 2014). Early endosomes mature into late endosomes with accumulation of these intraluminal vesicles (ILVs); upon fusing with the plasma membrane ILVs are released as exosomes into the extracellular environment (Colombo, Raposo & Thery, 2014). Due to their cone-shape structure, ceramide lipids induce negative curvature triggering inward budding of endosomal membranes (Trajkovic et al., 2008). We postulated that changes in lipid metabolism might therefore contribute to alterations in EV release by distinct oncogenes. We performed metabolomic analysis to determine how different oncogenes alter steady state abundance of lipid species. We discovered significant changes in different lipids from MCF10A cells transformed by CCND1, AURKB, MYC or HRAS relative to control MCF10A cells (**Figure 5A; Supplemental Table 4**). Many different lipid species were highly enriched in MYC- and HRAS-overexpressing cells compared to CCND1 and AURKB overexpressing MCF10A cells (**Figure 5A**).

Membrane curvature is one of the driving forces of plasma membrane and/or vesicle budding and can be affected by the lipid composition of membranes. For example, while lipids such as phosphatidylethanolamine (PE), phosphatic acid (PA), and diacylglycerol (DAG) which have smaller head groups favor negative curvature, lipids with large head groups such as lysophosphatidylcholine (LPC) or the large headgroups found in phosphatidylinositol phosphates (PI) support positive curvature (Zimmerberg, Kozlov, 2006, McMahon, Boucrot, 2015). MYC- and HRAS-overexpressing MCF10A cells were highly enriched with both LPC and PI lipids that favor positive membrane curvature **(Figure 5A)**. In addition, we found that ceramide lipids were significantly enriched only in MYC-overexpressing MCF10A cells **(Figure 5A)**. Ceramide can be generated via three major metabolic pathways as depicted in **Figure 5B**. The condensation of serine and fatty acyl-CoA initiates the *de novo* synthesis of ceramides. The breakdown of complex sphingolipids to sphingosine is the major source for the salvage pathway. Finally, ceramide is also produced through the hydrolysis of sphingomyelin to ceramide via the action of sphingomyelinases (SMases) (Kitatani, Idkowiak-Baldys & Hannun, 2008) (**Figure 5B**).

To evaluate lipid metabolic pathways that could contribute to EV release, we performed RNA sequencing of AURKB, MYC or HRAS overexpressing cells which we compared to control MCF10A cells. We sought to identify deregulated enzymes in the ceramide pathway. We found that many genes involved in ceramide production were dysregulated in all three oncogenic lines. However, N-SMases, SMPD3 and SMPD4, were significantly upregulated only in MYC overexpressing cells which have the greatest abundance of ceramide lipids **(Figure 5C)**.

These data suggest that MYC-overexpressing cells may use ceramide generation through sphingomyelin hydrolysis to produce high levels of exosomes. We tested the neutral sphingomyelinase inhibitor (GW4869) on a subset of oncogenic lines and measured EV release through nanoparticle tracking analysis. We observed a significant decrease in EV release from MYC and AURKB overexpressing cells, but not in control cells, or those expressing HRAS or CCND1 oncogenes **(Figure 5D)**. Interestingly, the MYC overexpressing cells were most responsive to GW4869 treatment, suggesting that ceramide-dependent EV production is especially important in MYC overexpressing cells.

We next asked if N-SMase inhibitor treatment can alter EV protein composition from cells transformed with different oncogenes **(Figure 5E)**. In MYC overexpressing cells, all vesicle protein markers tested exhibited a significant decrease in abundance **(Figure 5E)** with the exception of CD63 which showed variable expression **(Supplementary Figure 3)**. Similar to MYC EVs, the majority of the markers tested were decreased in AURKB EVs. However, an increase in levels of CD63 was observed in AURKB EVs upon inhibition of N-SMases **(Figure 5E** and **Supplementary Figure 3)**. In both AURKB and MYC EVs upon GW4869 treatment, *ALIX* was one of the most significantly downregulated proteins, supporting the role of N-SMases in the production of syndecan-syntenin-Alix containing exosomes (Baietti et al., 2012). However, we did not see a consistent decrease in CD63 levels as observed in Syndecan-Syntenin-Alix containing exosomes, suggesting that MYC and AURKB cells might produce other types of CD63 containing EVs from an alternative pathway upon N-SMase inhibition. Interestingly, we did not observe any significant change in the majority of the common small EV markers (e.g.; *CD9, CD63, ALIX, TSG101*) in HRAS EVs.

Furthermore, clathrin was significantly decreased in all three oncogenic EVs following GW4869 treatment, suggesting ceramide might be important in budding of early endosomal membranes originating from clathrin-mediated endocyctosis and/or clathrin coated vesicles budding from ER-Golgi network. Even though actinin-4 is a large EV marker (Kowal et al., 2016), it was significantly affected by N-SMase inhibition in HRAS EVs, strengthening the observation that ceramide might also play a role in budding from the plasma membranes (Menck et al., 2017).

In HRAS EVs we observe decreases in clathrin and actinin-4 abundance following GW4869 treatment but an increase in Annexin-V and CAV2 containing EVs **(Figure 5E** and **Supplementary Figure 3)**. These observations demonstrate that the ceramide pathway plays a major role in the biogenesis of small EVs from MYC high cells. In contrast, HRAS and CCND1 overexpressing cells release similar numbers of EVs upon N-SMase inhibition. For HRAS transformed cell derived EVs, while the total number of EVs was not significantly changed upon GW4869 treatment, we observed changes in the subtypes of released EVs.

### *TSG101* knockdown significantly decreases small EV release in AURKB and MYC overexpressing cells

The endosomal sorting complex required for transport (ESCRT) is a major regulator of MVB formation and is therefore also critical for EV release. ESCRT-0 complex recognizes and clusters ubiquitinated proteins, while ESCRT-I and -II complexes deform endosomal membranes to form buds where cargo is sorted. Finally, ESCRT-III together with accessory proteins cleave the buds to form intraluminal vesicles (ILVs) (Wollert, Hurley, 2010, Henne, Buchkovich & Emr, 2011, Colombo, Raposo & Thery, 2014) (model shown in **Figure 6A**). We sought to determine if ESCRT pathway components are altered in the context of specific oncogenes. We compared gene expression of ESCRT components in AURKB, MYC and HRAS overexpressing MCF10A cell extracts to control cell extracts. AURKB overexpressing cells exhibited the greatest number of dysregulated genes in the ESCRT pathway, whereas MYC showed the lowest numbered of altered genes compared to control cells (**Figure 6B**).

We next sought to determine if specific oncogene expressing cells are reliant on ESCRT components for EV release. We selected two ESCRT components previously identified to be important for EV release, *TSG101* and *ALIX* (also called *PDCP6IP*) (Colombo et al., 2013, Baietti et al., 2012), and depleted their expression via siRNAs. We then performed nanoparticle tracking analysis (NTA) on EVs from these depleted cells (**Figure 6C**). We did not observe significant changes in the number of released EVs in *TSG101* and *ALIX* knockdowns from control and HRAS-overexpressing MCF10A cells. However, *TSG101* knockdown significantly decreased the number of EVs from AURKB and MYC overexpressing cells, and this decrease was most pronounced in AURKB overexpressing cells (**Figure 6C**). Knocking down ALIX did not affect the numbers of EVs that were released. Thus, TSG101, but not ALIX, expression is important for mediating the number of EV release from MYC and AURKB expressing cells.

We next performed western blot analysis to determine if the protein content of released EVs changed following depletion of *TSG101* or *ALIX*. Amongst the 100K small EV fractions *TSG101* knockdown resulted in oncogene-specific alterations in EV protein expression. For example, in EVs from MYC high cells syntenin-1 protein was diminished, while clathrin was significantly up-regulated (**Figure 6E**). Although we did not observe any change in EV numbers from HRAS overexpressing cells upon *TSG101* knockdown **(Figure 6C)**, the levels of *CD81, ALIX* and syntenin-1 were significantly decreased from both 10K and 100K fractions demonstrating small EV population originating from endosomes decreased in these cells as well **(Figure 6D, 6E)**. However, EV pool seemed to be compensated by an increase in clathrin, *CAV2*, actinin-4, and Annexin-V containing EVs **(Figure 6D, 6E)**. Thus, TSG101 depletion resulted in diminished numbers of EVs produced from MYC and AURKB expressing cells **(Figure 6C)** and resulted in distinct alterations in protein content in EVs from all oncogenes tested **(Figure 6E)**.

*ALIX* depletion did not alter the number of released EVs in any of the three oncogenic lines tested **(Figure 6C)**. Nonetheless, *ALIX* knockdown diminished syntenin-1-containing vesicles in all oncogenes as expected (Baietti et al., 2012), and it also significantly increased the abundance of *CD81* and *TSG101* bearing vesicles especially from MYC- and AURKB-overexpressing cells **(Figure 6D, 6E)**, suggesting a mechanism for compensation of EV release. Upon *ALIX* knockdown, actinin-4 bearing EVs in 10K fraction were also significantly enriched in HRAS and MYC-derived EVs **(Figure 6D, 6E)**. Similarly, clathrin, Annexin-V and *RAB5* levels in 10K fraction were significantly higher in HRAS EVs **(Figure 6D,6E)**.

These results demonstrate that upon *TSG101* or *ALIX* depletion, oncogene-expressing MCF10A cells can induce EV release from alternative pathways (e.g.; either originating from MVBs or plasma membranes). This compensation seems to be oncogene-specific. Amongst the oncogenes tested, AURKB-overexpressing cells appear to rely on ESCRT-dependent pathways more than other oncogene-overexpressing cells. This is demonstrated by the reliance on *TSG101* expression by the AURKB-over expressing cells for EV release (**Figure 6C**).

### Downregulation of lysosome-regulated genes is correlated with increased EV release

Exosome secretion has been recently found to be regulated by ISG15 which causes ISGylation of *TSG101*, resulting in MVB colocalization with lysosomes and thus less exosome release (Villarroya-Beltri et al., 2016). Work by Villarroya-Beltri also found that lysosomal inhibition could increase exosome release in this context (Villarroya-Beltri et al., 2016). We explored if lysosomal function might be inhibited in MYC and AURKB expressing cells and if this might affect exosome secretion. We find that not only is ISG15 significantly downregulated, but also multiple lysosome-associated enzymes are downregulated as well in MYC and AURKB over expressing MCF10A cells (**Figure 7B**). Likewise, KEGG pathway analysis of RNA expression revealed that lysosomal function is one of the most downregulated pathways for MYC and AURKB overexpressing cells, but not HRAS high cells (**Figure 7A**). The MiT/TFE family of transcriptional factors, which include TFEB, TFE3 and MITF, are key regulators of lysosomal biogenesis and function (Perera, Zoncu, 2016). MiT/TFE factors are upregulated in pancreatic cancers and are required for maintaining high autophagy and lysosome function. The high levels of lysosome-mediated catabolic activity in pancreatic cancer is necessary to fuel bioenergetic needs of the tumors (Perera et al., 2015). Interestingly, TFE3 and TFEB are significantly downregulated in AURKB and MYC cells (**Figure 7B and 7C)**. These results are consistent with increased EV release from MYC and AURKB expressing cells. Thus, MYC and AURKB expressing cells might utilize high levels of EV secretion as an alternate mechanism for removing proteins or macromolecules that would otherwise be targeted for degradation via the lysosome.

**Figure 7.**
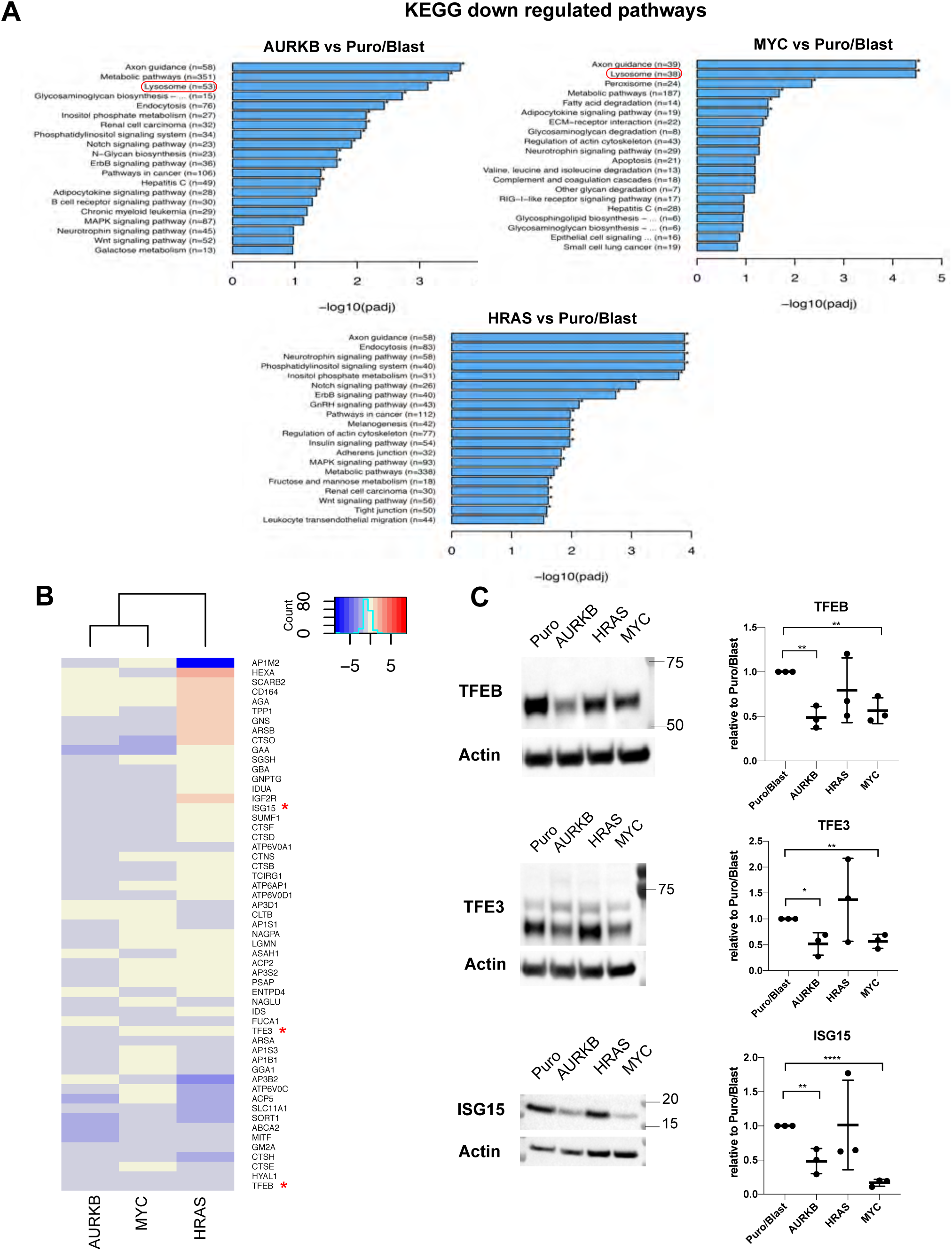
Downregulation of lysosome-associated genes is correlated with increased EV release. **A.** KEGG pathway analysis of down regulated pathways from AURKB, MYC and HRAS overexpressing cells compared to control (Puro/Blast). **B.** Heat-map of downregulated lysosome-associated genes from RNA-seq analysis of AURKB, MYC and HRAS overexpressing cells. **C.** Western blot analysis for *ISG15, TFEB* and *TFE3*. Represented images from three different experiment is on the left, the quantification of three different experiment is on the right side.

## Discussion

While oncogenes alter a variety of metabolic pathways to transform cells, they can also reprogram the size and content of released EVs to regulate the flux of cellular biomass and/or to enhance intercellular communication. In the case of MYC overexpressing cells, we find that up to 7% of cellular volume may be shed via EVs every two days, at least 20 times more than control cells and several fold more than those transformed by other oncogenes. MYC significantly alters ceramide metabolism by upregulating sphingomyelinases and this results in high levels of ceramides, which favor the accumulation of small EVs that originate from endosomal membranes. AURKB-overexpressing cells, in contrast, are highly dependent on the ESCRT pathway. While HRAS (G12V) expression strongly favors the release of actinin-4, flotillin-1, and Annexin-V-bearing EVs that might originate from plasma membranes. Thus, our studies reveal that distinct oncogenes modify the number, cargo and the likely origin of released EVs from cancer cells.

Our work strengthens the hypothesis that oncogenes direct the release of large numbers of EVs (Al-Nedawi et al., 2008, Balaj et al., 2011, Bebelman et al., 2018, Choi et al., 2017), and demonstrates that the degree of EV production depends on the driver oncogene. MYC and AURKB overexpressing cells release a disproportionately high number of small EVs. MYC recapitulated the increase in EV production in two other cell types tested, suggesting that MYC-dependent increase in EV release could be a common feature of increased biomass flux and may thus represent a new metabolic parameter to consider.

Furthermore, EV populations are heterogeneous and complex. Recently, several distinct subtypes of EVs have emerged from studies performing proteomic analysis after immunoprecipitation of specific surface markers and high-resolution density gradient fractionation (Jeppesen et al., 2019, Coulter et al., 2018, Kowal et al., 2016). However, the field still lacks clear methods to distinguish various populations of EVs since similar size vesicles could be either microvesicles or exosomes, and same types of tetraspanins could be presented on vesicles of endosomal or plasma membrane origin. Additionally, both lipids and ESCRT components are involved in budding of membranes from either plasma membranes or MVBs (Mathieu et al., 2019). Our studies show that although they may share some common characteristics, each oncogene-expressing EV population has its unique composition. MYC and AURKB EVs share similar exosome markers but *RAB5* is enriched in AURKB-derived EVs whereas clathrin-containing EVs are highly enriched in MYC-derived EVs. However, HRAS EVs are distinct form other oncogene-derived EVs and highly enriched with large EV markers like actinin-4, and flotillin-1. To our knowledge, our data is the first to describe how various oncogenes might alter heterogeneity of released EVs. Future studies should further dissect the subpopulations of these EVs using high-density gradient fractionation and immunoisolation to identify unique markers of oncogene specific EVs that may be useful as specific biomarker and potential therapeutics.

In addition to the heterogeneity of protein composition observed, we also found that miRNAs may be released in EVs or retained in cells in an oncogene-dependent manner. MYC and AURKB oncogenes share many common miRNAs enriched in EVs, whereas the miRNA content of CCND1 EVs is similar to EVs released from control cells. We also identified uniquely enriched miRNAs loaded into EVs for a given oncogene. Our analysis could be further extended to identify oncogene specific EV RNA biomarkers for different cancer types.

In the present study, we explored known miRNA motifs that have been identified to affect miRNA sorting into EVs. Both hnRNPA2B1 and SYNCRIP binding motifs are present in miRNAs loaded into oncogene derived MCF10A EVs; the hnRNPA2B1 motif was more abundant in all oncogene-derived EVs tested. Moreover, these two motifs were only detected in half of the EV enriched miRNAs. Thus, perhaps other RNA motifs or sorting principals may exist to regulate the sorting of other miRNAs into EVs. We sought to identify unknown motifs unique to a given oncogene-derived EV. We used Improbizer to identify common motif sequences in miRNAs only enriched in MYC EVs, but we could not identify statistically significant E-values for these sequences from MEME analysis (Ao et al., 2004, Bailey et al., 2009). This was probably due to the small number of unique miRNAs (<10 miRNAs) enriched in MYC EVs (and similarly for other oncogenes). In future studies, high-throughput small RNA-sequencing and proteomic analysis of oncogene-derived MCF10A EVs may identify additional RNA binding proteins and miRNA motifs that could have roles in EV miRNA sorting in an oncogene-specific manner.

In the present study we find that specific oncogenes reprogram gene expression profiles of EV biogenesis pathways. The MYC oncogene significantly alters ceramide lipid metabolism that has been previously shown to trigger budding of endosomal membranes to produce exosomes. The blocking of this pathway via N-SMase inhibition significantly affected MYC dependent EV release. With N-SMase inhibition, we observed a significant decrease in *ALIX*, clathrin, and *RAB5* containing vesicles both in AURKB and MYC EV populations, suggesting that these vesicles are coming from endosomal origin. Even though AURKB EVs were diminished following N-SMase inhibition, the decrease was not as pronounced as in MYC EVs. The accumulation of *CD63* and *TSG101* bearing vesicles in AURKB EVs treated with N-SMase suggest that exosomes produced via ESCRT-dependent pathways can partially compensating for EV release in AURKB-overexpressing MCF10A cells (Figure 5E and Supplementary Figure 3).

Previous studies have shown the importance of different components of the ESCRT machinery in the biogenesis of EVs (Colombo et al., 2013, Baietti et al., 2012, Coulter et al., 2018). Our study not only strengthened these observations, but also provides strong evidence for additional complexity and heterogeneity in the EV pool. The compensation of EV production via alternative types of vesicles became evident when ESCRT pathway components were silenced via siRNA treatment. *TSG101* knockdown significantly reduced EV production in AURKB and MYC cells. *CD81* was highly correlated with *TSG101* bearing vesicles since *TSG101* knockdown significantly reduced *CD81* in all oncogenes tested, but also *ALIX* knockdown significantly increased both *CD81* and *TSG101* bearing vesicles in MYC and AURKB EVs suggesting *CD81* and *TSG101* originate from the same ESCRT-dependent pathway. In contrast, *TSG101* knockdown affects syntenin-1 and *ALIX* containing EVs in HRAS and MYC EVs, reinforcing the role of *TSG101* in the biogenesis of Syntenin-ALIX containing vesicles (Baietti et al., 2012). Interestingly, clathrin is significantly upregulated upon TSG101 depletion in MYC derived EVs whereas it is significantly downregulated during N-SMase inhibition. Thus, ceramide pathway might partially compensate for released EVs from MYC cells when TSG101 depleted.

Oncogenes reprogram tumor cell metabolism by upregulating pathways such as glycolysis, glutaminolysis and increased lipid synthesis. These anabolic processes contribute to increases in cellular biomass which in turn permit accelerated cell division. However, to maintain cellular biomass homeostasis tumor cells need to also degrade excess macromolecules or alternately release them via extracellular vesicles. We find that many oncogenes, but especially MYC and AURKB, can increase EV release from tumor cells. Through increased EV release, tumor cells may thus balance anabolic processes by jettisoning unwanted macromolecules, such as tumor suppressive miRNAs, toxic lipids and other cargos. Here we propose that understanding the principals that guide regulated release of extracellular vesicles in cancer cells represents a fundamentally new way that oncogenes reprogram cellular metabolism.

## Acknowledgements

This publication is part of the NIH Extracellular RNA Communication Consortium paper package and was supported by the NIH Common Fund’s exRNA Communication Program. We thank Andrew Leidal and Jayanta Debnath for equipment and reagent sharing. We thank all the members of Goga lab for their help and support. We thank the staff at the University of California Berkeley Electron Microscope Laboratory for advice and assistance in electron microscopy sample preparation and data collection. The work was funded by NIH (1U19CA179512 to A.G. and N.D.L.), (R01CA223817 to A.G. and R01 DC005991 to N.D.L) and S10RR026758. The CDMRP (W81XWH-18-1-0713 to A.G.) and the Gazarian Family Endowment (to A.G.). Also, the US NIH F99/K00 Predoctoral to Postdoctoral Transition Award (F99CA212488 to R.C.). R.M.P is the Nadia’s Gift Foundation Innovator of the Damon Runyon Cancer Research Foundation (DRR-46-17) and is supported by an NIH Director’s New Innovator Award (1DP2CA216364) and Pancreatic Cancer Action Network Career Development Award.

---

**Supplementary Figure 1.**
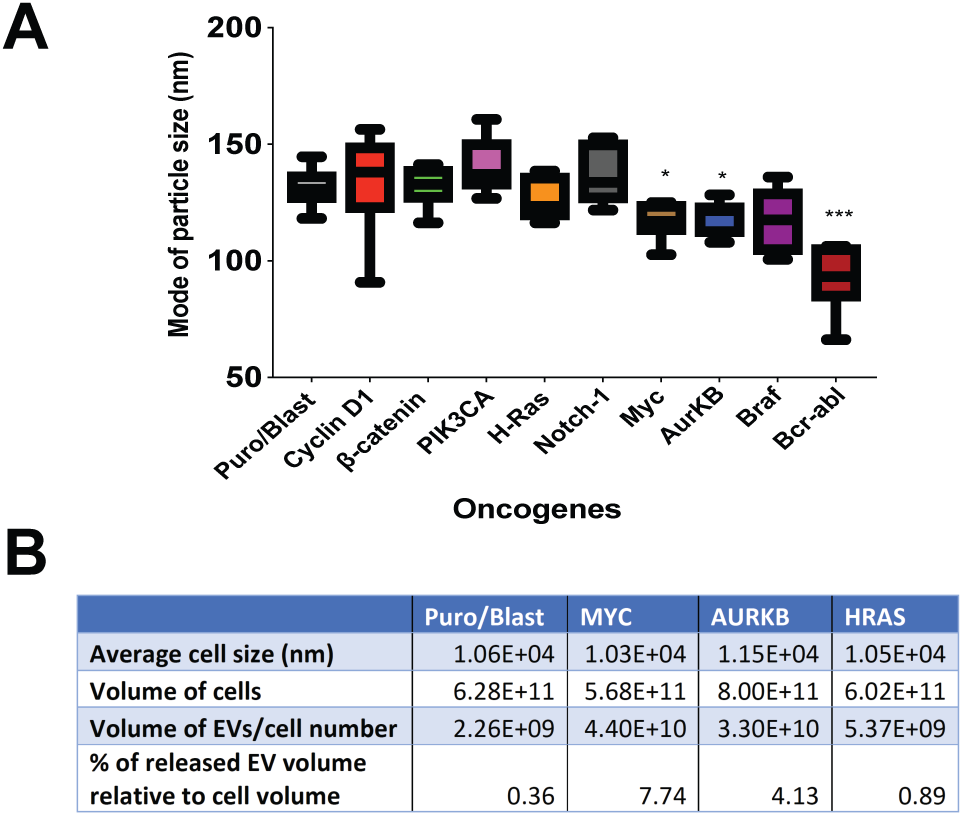
Related to Figure 1. Size and volume analysis of EVs from MCF10A oncogenic lines. **A**.Mode of particle sizes from Nanoparticle tracking analysis (NTA) (from Figure 1C and 1D). Box and whisker blots from 6 different NTA are plotted. *p<0.05, ***p<0.001.**B.** Cell volume and released EV volume for each cell line are presented.

**Supplementary Figure 2.**
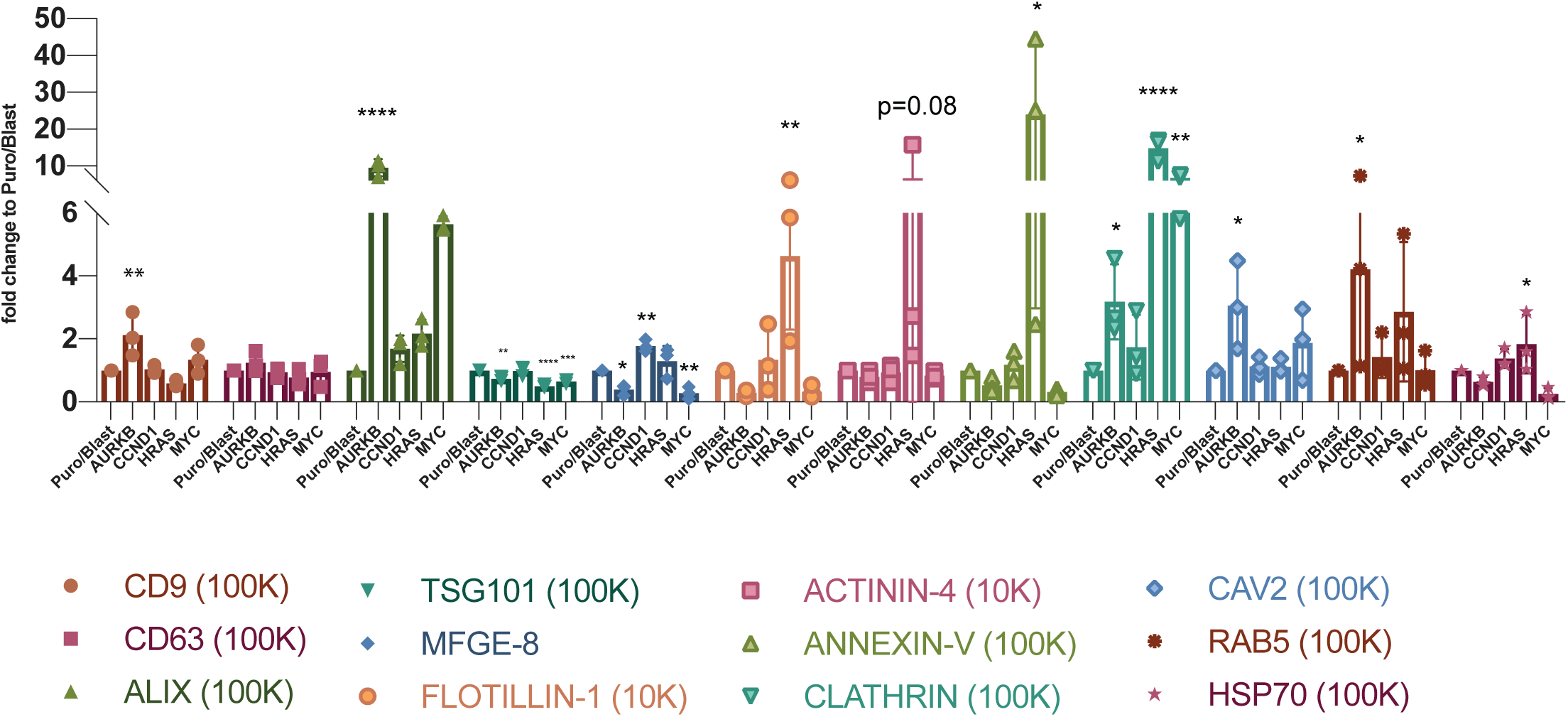
Related to Figure 3. Quantification of EV marker analysis. Three independent experiments are performed, and quantification is performed by ImageLab software (BIORAD). Each marker is normalized to control (Puro/Blast) in each group and represented as bar graphs.

**Supplementary Figure 3.**
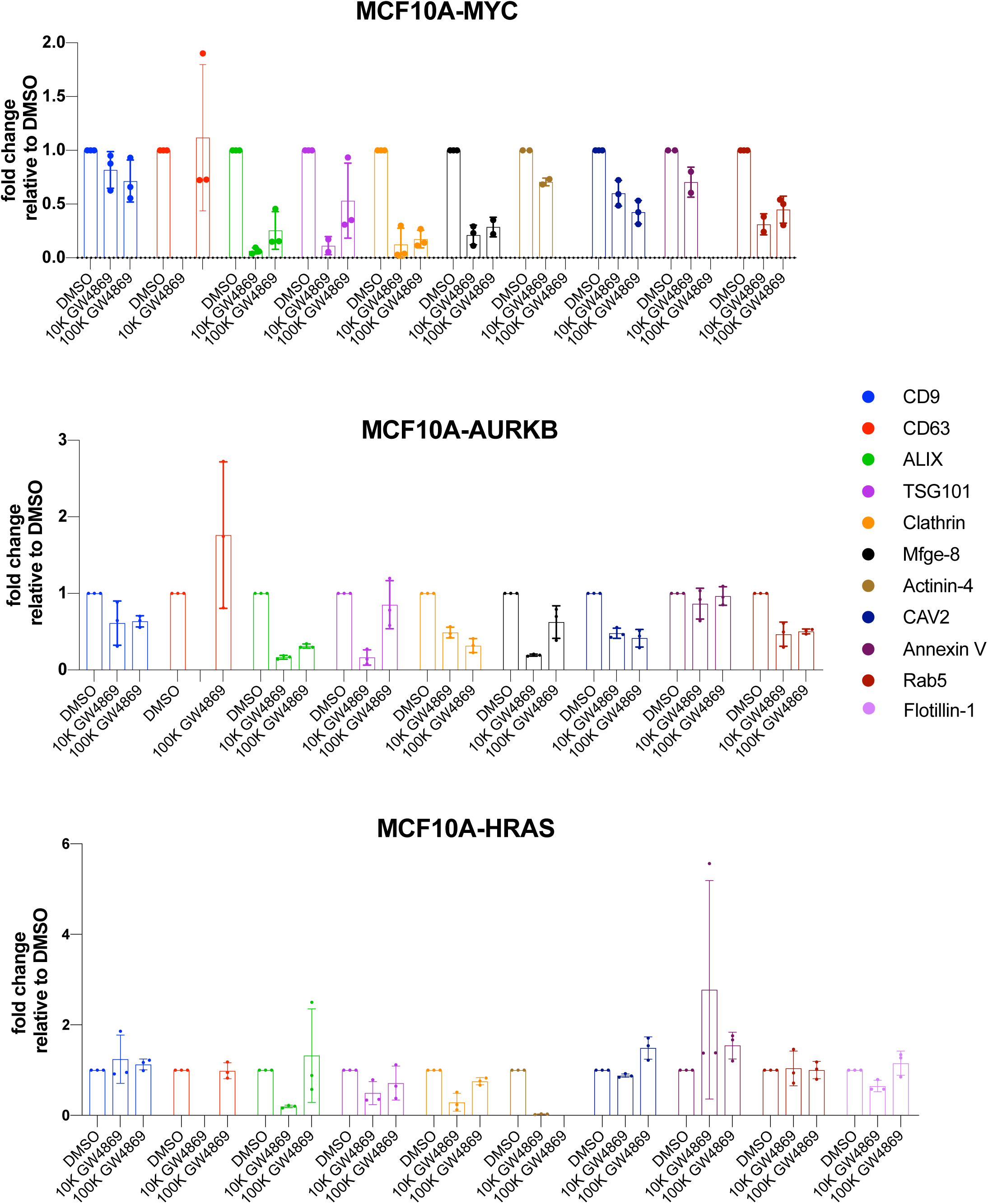
Quantification of western blot analysis from MCF10A cells treated either with vehicle (DMSO) or N-SMase inhibitor, GW4869 (SµM), Related to Figure 5. Three independent experiments performed for all oncogenes tested. Quantification of protein bands is performed using ImageLab software from BioRAD. Fold change of EV markers relative to DMSO treatment in each group compared and values are depicted in the bar graph.

**Supplementary Table 1.**
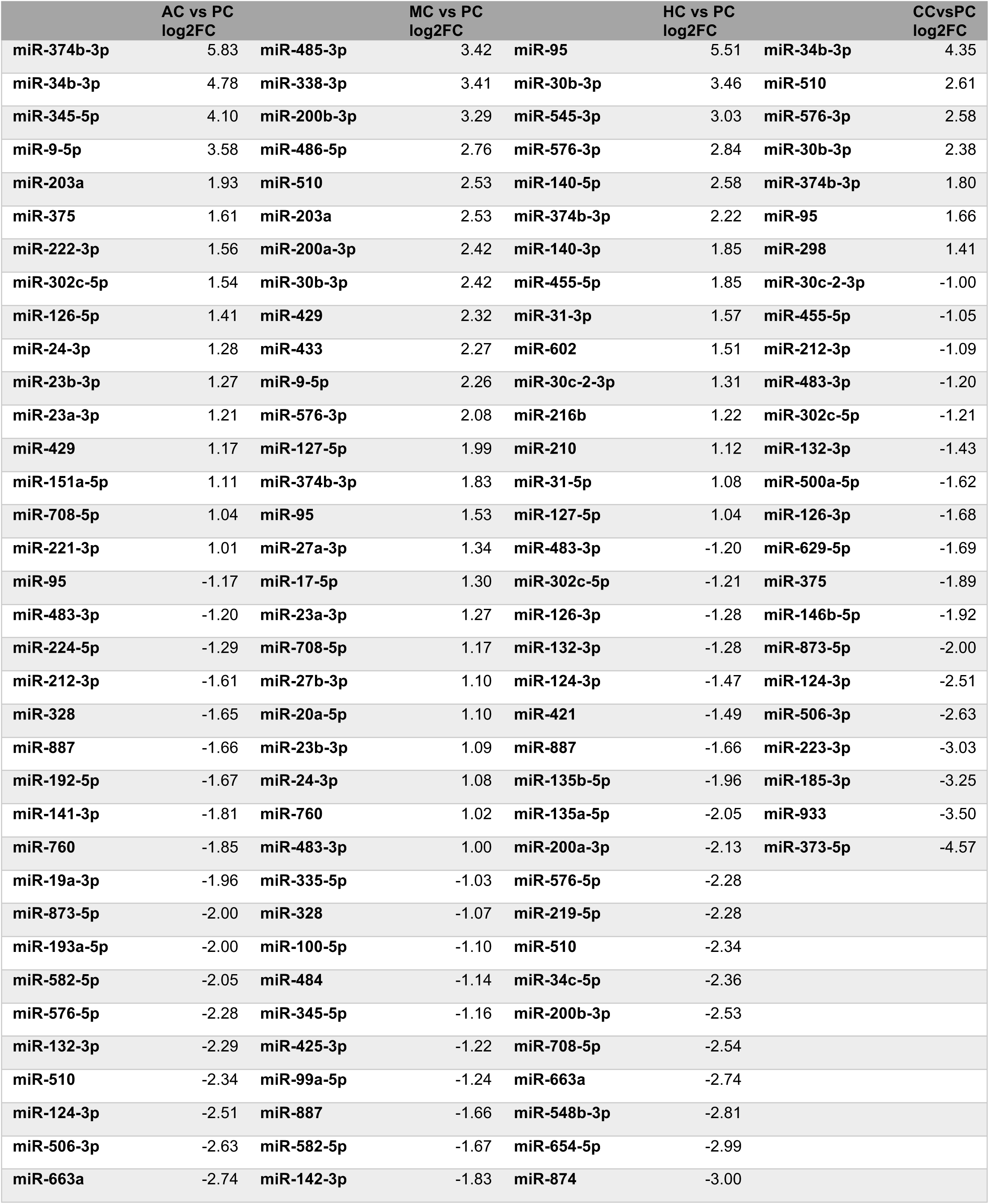
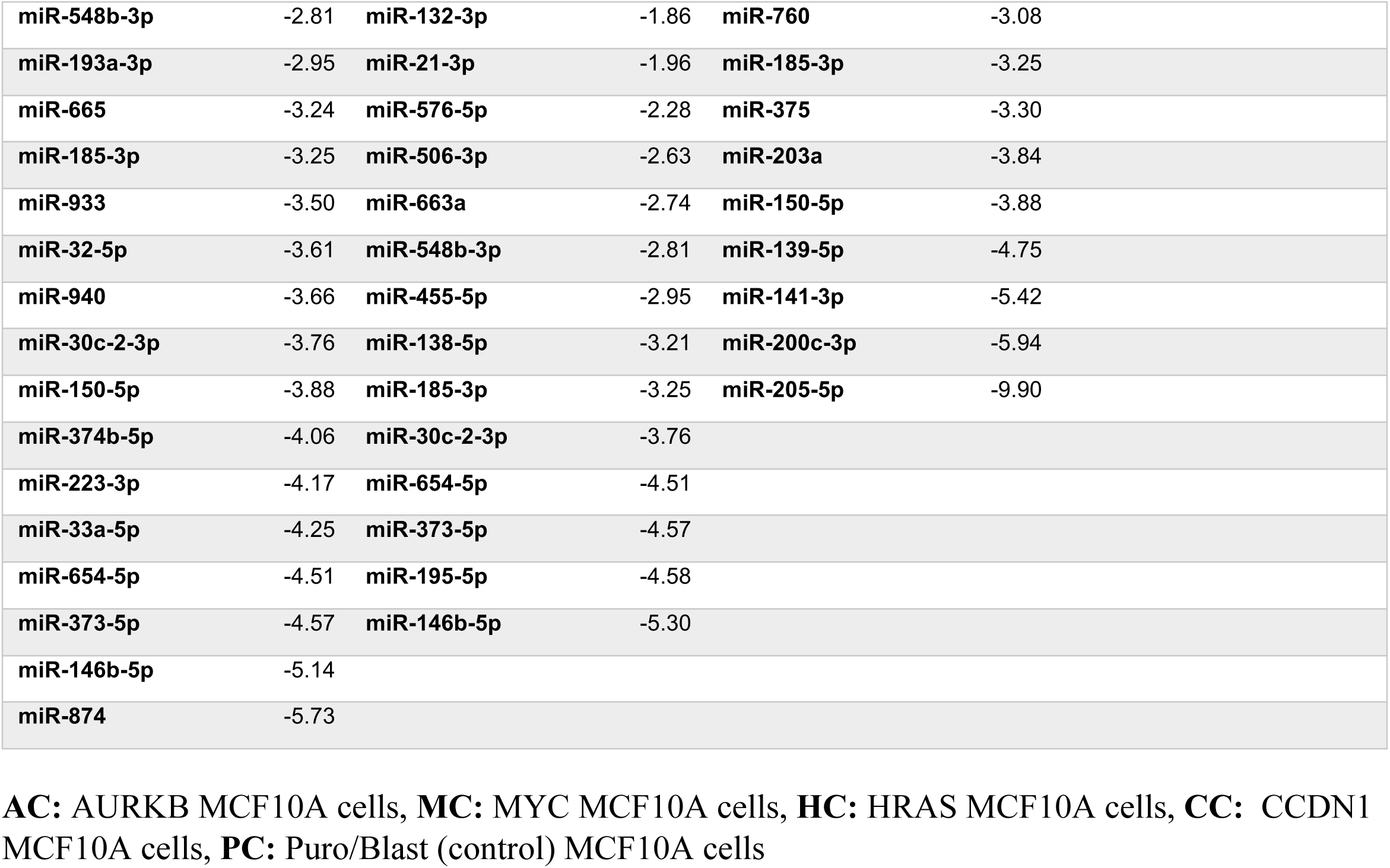
miRNAs differentially regulated in cells for a given oncogene compared to control MCF10A cells, Related to Figure 4.

|  | AC vs PC<br>log2FC |  | MC vs PC<br>log2FC |  | HC vs PC<br>log2FC |  | CCvsPC<br>log2FC |
| --- | --- | --- | --- | --- | --- | --- | --- |
| miR-374b-3p | 5.83 | miR-485-3p | 3.42 | miR-95 | 5.51 | miR-34b-3p | 4.35 |
| miR-34b-3p | 4.78 | miR-338-3p | 3.41 | miR-30b-3p | 3.46 | miR-510 | 2.61 |
| miR-345-5p | 4.10 | miR-200b-3p | 3.29 | miR-545-3p | 3.03 | miR-576-3p | 2.58 |
| miR-9-5p | 3.58 | miR-486-5p | 2.76 | miR-576-3p | 2.84 | miR-30b-3p | 2.38 |
| miR-203a | 1.93 | miR-510 | 2.53 | miR-140-5p | 2.58 | miR-374b-3p | 1.80 |
| miR-375 | 1.61 | miR-203a | 2.53 | miR-374b-3p | 2.22 | miR-95 | 1.66 |
| miR-222-3p | 1.56 | miR-200a-3p | 2.42 | miR-140-3p | 1.85 | miR-298 | 1.41 |
| miR-302c-5p | 1.54 | miR-30b-3p | 2.42 | miR-455-5p | 1.85 | miR-30c-2-3p | -1.00 |
| miR-126-5p | 1.41 | miR-429 | 2.32 | miR-31-3p | 1.57 | miR-455-5p | -1.05 |
| miR-24-3p | 1.28 | miR-433 | 2.27 | miR-602 | 1.51 | miR-212-3p | -1.09 |
| miR-23b-3p | 1.27 | miR-9-5p | 2.26 | miR-30c-2-3p | 1.31 | miR-483-3p | -1.20 |
| miR-23a-3p | 1.21 | miR-576-3p | 2.08 | miR-216b | 1.22 | miR-302c-5p | -1.21 |
| miR-429 | 1.17 | miR-127-5p | 1.99 | miR-210 | 1.12 | miR-132-3p | -1.43 |
| miR-151a-5p | 1.11 | miR-374b-3p | 1.83 | miR-31-5p | 1.08 | miR-500a-5p | -1.62 |
| miR-708-5p | 1.04 | miR-95 | 1.53 | miR-127-5p | 1.04 | miR-126-3p | -1.68 |
| miR-221-3p | 1.01 | miR-27a-3p | 1.34 | miR-483-3p | -1.20 | miR-629-5p | -1.69 |
| miR-95 | -1.17 | miR-17-5p | 1.30 | miR-302c-5p | -1.21 | miR-375 | -1.89 |
| miR-483-3p | -1.20 | miR-23a-3p | 1.27 | miR-126-3p | -1.28 | miR-146b-5p | -1.92 |
| miR-224-5p | -1.29 | miR-708-5p | 1.17 | miR-132-3p | -1.28 | miR-873-5p | -2.00 |
| miR-212-3p | -1.61 | miR-27b-3p | 1.10 | miR-124-3p | -1.47 | miR-124-3p | -2.51 |
| miR-328 | -1.65 | miR-20a-5p | 1.10 | miR-421 | -1.49 | miR-506-3p | -2.63 |
| miR-887 | -1.66 | miR-23b-3p | 1.09 | miR-887 | -1.66 | miR-223-3p | -3.03 |
| miR-192-5p | -1.67 | miR-24-3p | 1.08 | miR-135b-5p | -1.96 | miR-185-3p | -3.25 |
| miR-141-3p | -1.81 | miR-760 | 1.02 | miR-135a-5p | -2.05 | miR-933 | -3.50 |
| miR-760 | -1.85 | miR-483-3p | 1.00 | miR-200a-3p | -2.13 | miR-373-5p | -4.57 |
| miR-19a-3p | -1.96 | miR-335-5p | -1.03 | miR-576-5p | -2.28 |  |  |
| miR-873-5p | -2.00 | miR-328 | -1.07 | miR-219-5p | -2.28 |  |  |
| miR-193a-5p | -2.00 | miR-100-5p | -1.10 | miR-510 | -2.34 |  |  |
| miR-582-5p | -2.05 | miR-484 | -1.14 | miR-34c-5p | -2.36 |  |  |
| miR-576-5p | -2.28 | miR-345-5p | -1.16 | miR-200b-3p | -2.53 |  |  |
| miR-132-3p | -2.29 | miR-425-3p | -1.22 | miR-708-5p | -2.54 |  |  |
| miR-510 | -2.34 | miR-99a-5p | -1.24 | miR-663a | -2.74 |  |  |
| miR-124-3p | -2.51 | miR-887 | -1.66 | miR-548b-3p | -2.81 |  |  |
| miR-506-3p | -2.63 | miR-582-5p | -1.67 | miR-654-5p | -2.99 |  |  |
| miR-663a | -2.74 | miR-142-3p | -1.83 | miR-874 | -3.00 |  |  |

|  |  |  |  |  |  |
| --- | --- | --- | --- | --- | --- |
| **miR-548b-3p** | -2.81 | **miR-132-3p** | -1.86 | **miR-760** | -3.08 |
| **miR-193a-3p** | -2.95 | **miR-21-3p** | -1.96 | **miR-185-3p** | -3.25 |
| **miR-665** | -3.24 | **miR-576-5p** | -2.28 | **miR-375** | -3.30 |
| **miR-185-3p** | -3.25 | **miR-506-3p** | -2.63 | **miR-203a** | -3.84 |
| **miR-933** | -3.50 | **miR-663a** | -2.74 | **miR-150-5p** | -3.88 |
| **miR-32-5p** | -3.61 | **miR-548b-3p** | -2.81 | **miR-139-5p** | -4.75 |
| **miR-940** | -3.66 | **miR-455-5p** | -2.95 | **miR-141-3p** | -5.42 |
| **miR-30c-2-3p** | -3.76 | **miR-138-5p** | -3.21 | **miR-200c-3p** | -5.94 |
| **miR-150-5p** | -3.88 | **miR-185-3p** | -3.25 | **miR-205-5p** | -9.90 |
| **miR-374b-5p** | -4.06 | **miR-30c-2-3p** | -3.76 |  |  |
| **miR-223-3p** | -4.17 | **miR-654-5p** | -4.51 |  |  |
| **miR-33a-5p** | -4.25 | **miR-373-5p** | -4.57 |  |  |
| **miR-654-5p** | -4.51 | **miR-195-5p** | -4.58 |  |  |
| **miR-373-5p** | -4.57 | **miR-146b-5p** | -5.30 |  |  |
| **miR-146b-5p** | -5.14 |  |  |  |  |
| **miR-874** | -5.73 |  |  |  |  |

**AC:** AURKB MCF10A cells, **MC:** MYC MCF10A cells, **HC:** HRAS MCF10A cells, **CC:** CCDN1 MCF10A cells, **PC:** Puro/Blast (control) MCF10A cells

**Supplementary Table 2.**
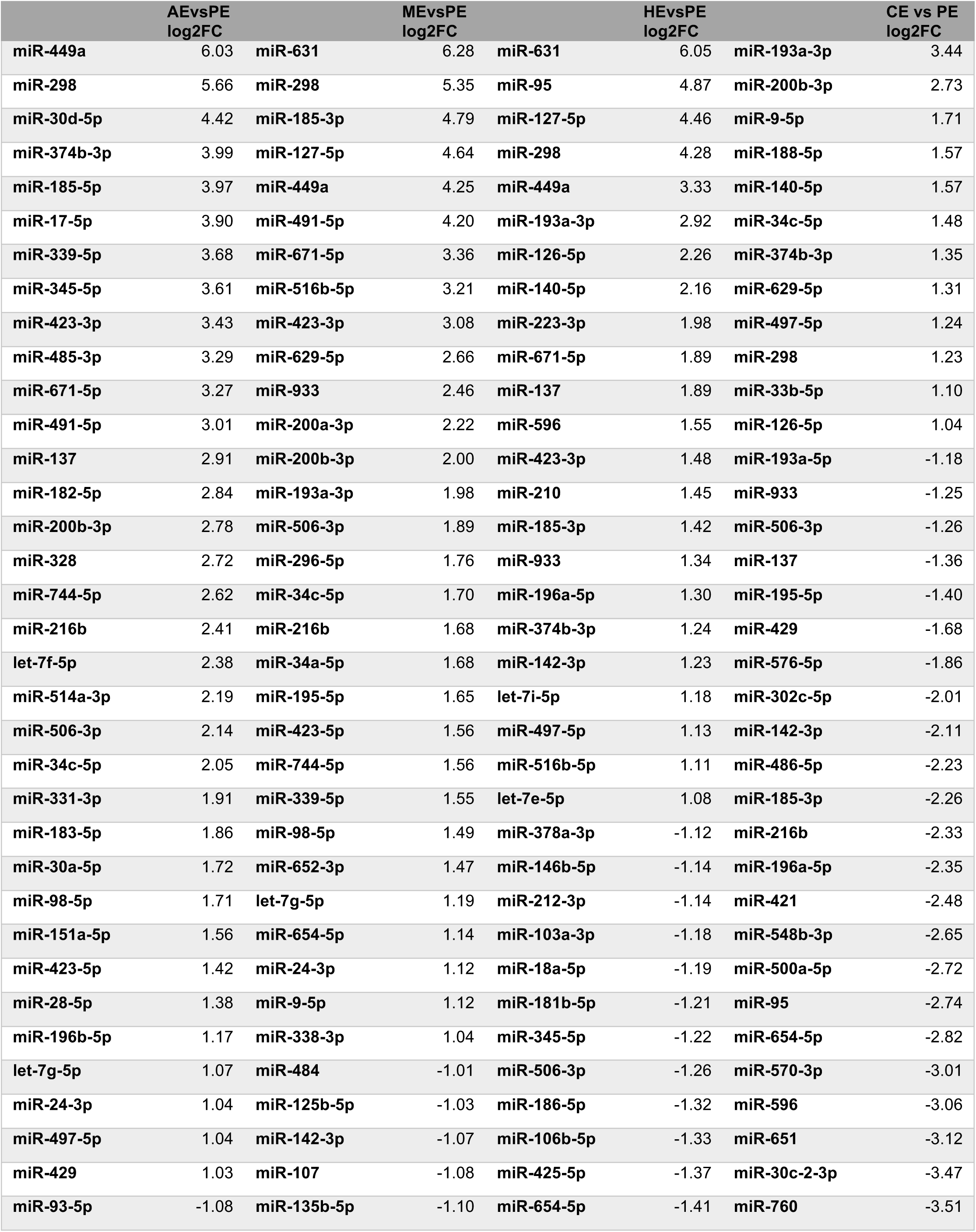
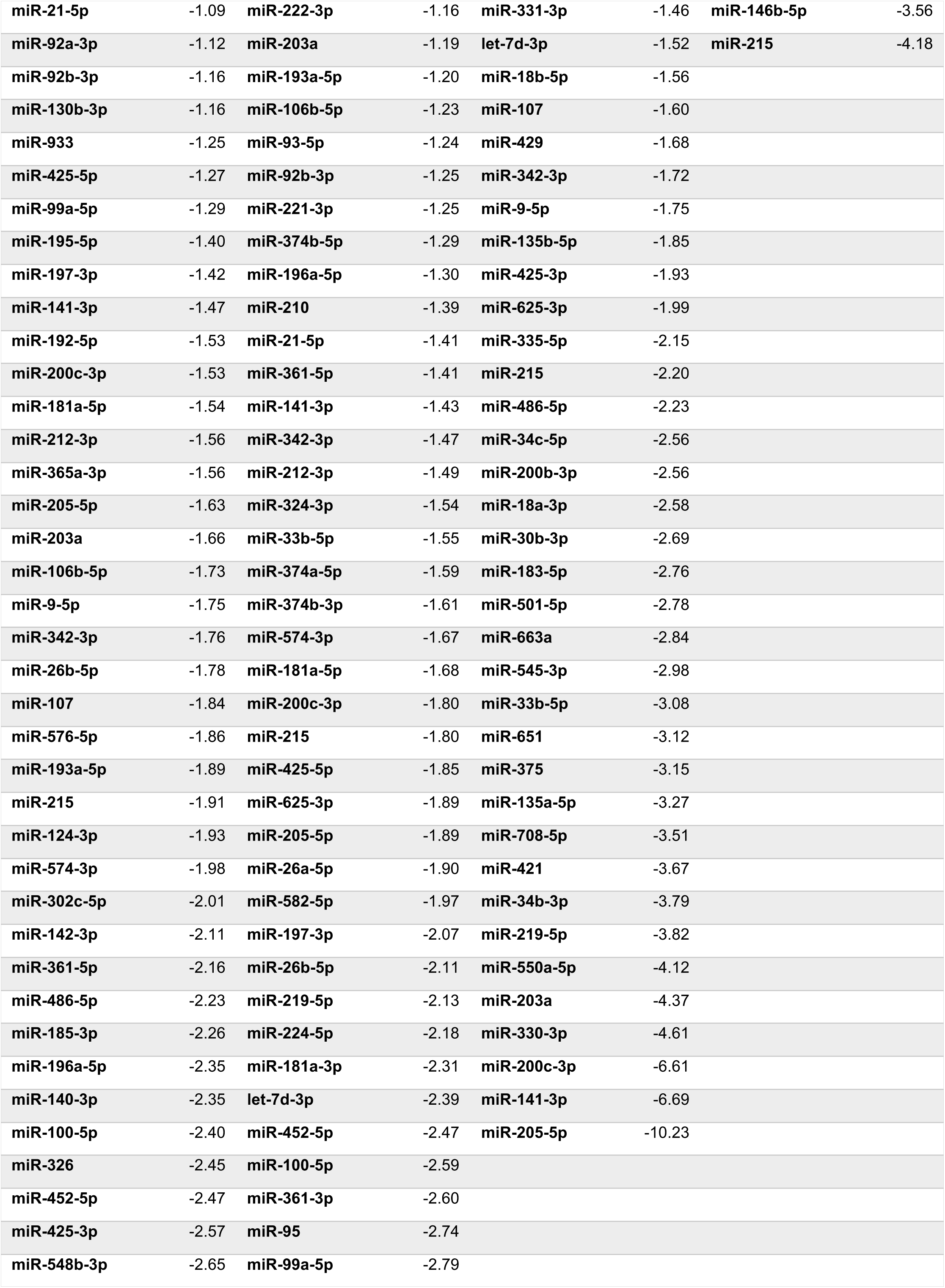
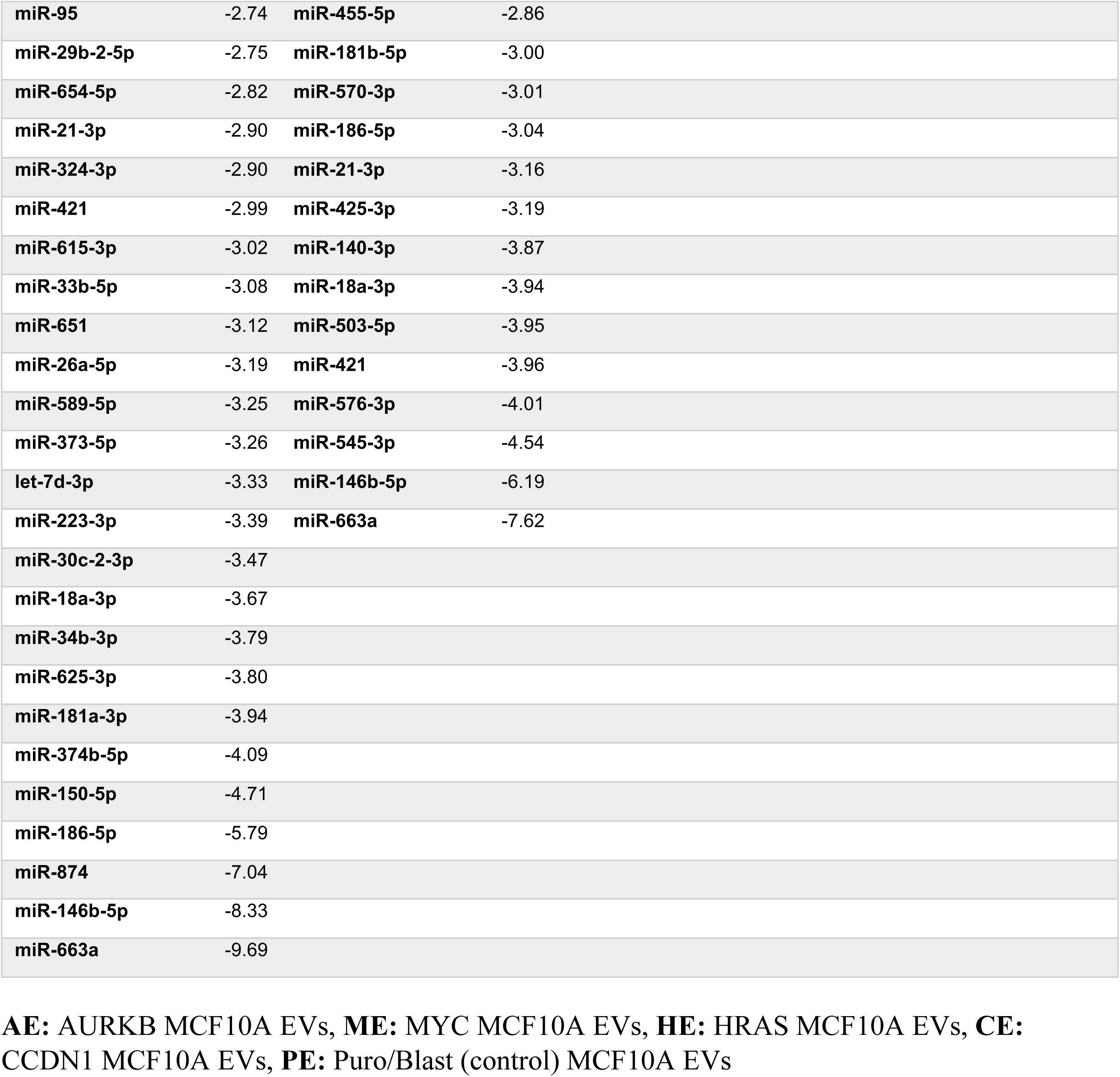
miRNAs differentially sorted into EVs for a given oncogene compared to control MCF10A EVs, Related to Figure 4.

|  | AEvsPE<br>log2FC |  | MEvsPE<br>log2FC |  | HEvsPE<br>log2FC |  | CE vs PE<br>log2FC |
| --- | --- | --- | --- | --- | --- | --- | --- |
| miR-449a | 6.03 | miR-631 | 6.28 | miR-631 | 6.05 | miR-193a-3p | 3.44 |
| miR-298 | 5.66 | miR-298 | 5.35 | miR-95 | 4.87 | miR-200b-3p | 2.73 |
| miR-30d-5p | 4.42 | miR-185-3p | 4.79 | miR-127-5p | 4.46 | miR-9-5p | 1.71 |
| miR-374b-3p | 3.99 | miR-127-5p | 4.64 | miR-298 | 4.28 | miR-188-5p | 1.57 |
| miR-185-5p | 3.97 | miR-449a | 4.25 | miR-449a | 3.33 | miR-140-5p | 1.57 |
| miR-17-5p | 3.90 | miR-491-5p | 4.20 | miR-193a-3p | 2.92 | miR-34c-5p | 1.48 |
| miR-339-5p | 3.68 | miR-671-5p | 3.36 | miR-126-5p | 2.26 | miR-374b-3p | 1.35 |
| miR-345-5p | 3.61 | miR-516b-5p | 3.21 | miR-140-5p | 2.16 | miR-629-5p | 1.31 |
| miR-423-3p | 3.43 | miR-423-3p | 3.08 | miR-223-3p | 1.98 | miR-497-5p | 1.24 |
| miR-485-3p | 3.29 | miR-629-5p | 2.66 | miR-671-5p | 1.89 | miR-298 | 1.23 |
| miR-671-5p | 3.27 | miR-933 | 2.46 | miR-137 | 1.89 | miR-33b-5p | 1.10 |
| miR-491-5p | 3.01 | miR-200a-3p | 2.22 | miR-596 | 1.55 | miR-126-5p | 1.04 |
| miR-137 | 2.91 | miR-200b-3p | 2.00 | miR-423-3p | 1.48 | miR-193a-5p | -1.18 |
| miR-182-5p | 2.84 | miR-193a-3p | 1.98 | miR-210 | 1.45 | miR-933 | -1.25 |
| miR-200b-3p | 2.78 | miR-506-3p | 1.89 | miR-185-3p | 1.42 | miR-506-3p | -1.26 |
| miR-328 | 2.72 | miR-296-5p | 1.76 | miR-933 | 1.34 | miR-137 | -1.36 |
| miR-744-5p | 2.62 | miR-34c-5p | 1.70 | miR-196a-5p | 1.30 | miR-195-5p | -1.40 |
| miR-216b | 2.41 | miR-216b | 1.68 | miR-374b-3p | 1.24 | miR-429 | -1.68 |
| let-7f-5p | 2.38 | miR-34a-5p | 1.68 | miR-142-3p | 1.23 | miR-576-5p | -1.86 |
| miR-514a-3p | 2.19 | miR-195-5p | 1.65 | let-7i-5p | 1.18 | miR-302c-5p | -2.01 |
| miR-506-3p | 2.14 | miR-423-5p | 1.56 | miR-497-5p | 1.13 | miR-142-3p | -2.11 |
| miR-34c-5p | 2.05 | miR-744-5p | 1.56 | miR-516b-5p | 1.11 | miR-486-5p | -2.23 |
| miR-331-3p | 1.91 | miR-339-5p | 1.55 | let-7e-5p | 1.08 | miR-185-3p | -2.26 |
| miR-183-5p | 1.86 | miR-98-5p | 1.49 | miR-378a-3p | -1.12 | miR-216b | -2.33 |
| miR-30a-5p | 1.72 | miR-652-3p | 1.47 | miR-146b-5p | -1.14 | miR-196a-5p | -2.35 |
| miR-98-5p | 1.71 | let-7g-5p | 1.19 | miR-212-3p | -1.14 | miR-421 | -2.48 |
| miR-151a-5p | 1.56 | miR-654-5p | 1.14 | miR-103a-3p | -1.18 | miR-548b-3p | -2.65 |
| miR-423-5p | 1.42 | miR-24-3p | 1.12 | miR-18a-5p | -1.19 | miR-500a-5p | -2.72 |
| miR-28-5p | 1.38 | miR-9-5p | 1.12 | miR-181b-5p | -1.21 | miR-95 | -2.74 |
| miR-196b-5p | 1.17 | miR-338-3p | 1.04 | miR-345-5p | -1.22 | miR-654-5p | -2.82 |
| let-7g-5p | 1.07 | miR-484 | -1.01 | miR-506-3p | -1.26 | miR-570-3p | -3.01 |
| miR-24-3p | 1.04 | miR-125b-5p | -1.03 | miR-186-5p | -1.32 | miR-596 | -3.06 |
| miR-497-5p | 1.04 | miR-142-3p | -1.07 | miR-106b-5p | -1.33 | miR-651 | -3.12 |
| miR-429 | 1.03 | miR-107 | -1.08 | miR-425-5p | -1.37 | miR-30c-2-3p | -3.47 |
| miR-93-5p | -1.08 | miR-135b-5p | -1.10 | miR-654-5p | -1.41 | miR-760 | -3.51 |
| miR-21-5p | -1.09 | miR-222-3p | -1.16 | miR-331-3p | -1.46 | miR-146b-5p | -3.56 |
| miR-92a-3p | -1.12 | miR-203a | -1.19 | let-7d-3p | -1.52 | miR-215 | -4.18 |
| miR-92b-3p | -1.16 | miR-193a-5p | -1.20 | miR-18b-5p | -1.56 |  |  |
| miR-130b-3p | -1.16 | miR-106b-5p | -1.23 | miR-107 | -1.60 |  |  |
| miR-933 | -1.25 | miR-93-5p | -1.24 | miR-429 | -1.68 |  |  |
| miR-425-5p | -1.27 | miR-92b-3p | -1.25 | miR-342-3p | -1.72 |  |  |
| miR-99a-5p | -1.29 | miR-221-3p | -1.25 | miR-9-5p | -1.75 |  |  |
| miR-195-5p | -1.40 | miR-374b-5p | -1.29 | miR-135b-5p | -1.85 |  |  |
| miR-197-3p | -1.42 | miR-196a-5p | -1.30 | miR-425-3p | -1.93 |  |  |
| miR-141-3p | -1.47 | miR-210 | -1.39 | miR-625-3p | -1.99 |  |  |
| miR-192-5p | -1.53 | miR-21-5p | -1.41 | miR-335-5p | -2.15 |  |  |
| miR-200c-3p | -1.53 | miR-361-5p | -1.41 | miR-215 | -2.20 |  |  |
| miR-181a-5p | -1.54 | miR-141-3p | -1.43 | miR-486-5p | -2.23 |  |  |
| miR-212-3p | -1.56 | miR-342-3p | -1.47 | miR-34c-5p | -2.56 |  |  |
| miR-365a-3p | -1.56 | miR-212-3p | -1.49 | miR-200b-3p | -2.56 |  |  |
| miR-205-5p | -1.63 | miR-324-3p | -1.54 | miR-18a-3p | -2.58 |  |  |
| miR-203a | -1.66 | miR-33b-5p | -1.55 | miR-30b-3p | -2.69 |  |  |
| miR-106b-5p | -1.73 | miR-374a-5p | -1.59 | miR-183-5p | -2.76 |  |  |
| miR-9-5p | -1.75 | miR-374b-3p | -1.61 | miR-501-5p | -2.78 |  |  |
| miR-342-3p | -1.76 | miR-574-3p | -1.67 | miR-663a | -2.84 |  |  |
| miR-26b-5p | -1.78 | miR-181a-5p | -1.68 | miR-545-3p | -2.98 |  |  |
| miR-107 | -1.84 | miR-200c-3p | -1.80 | miR-33b-5p | -3.08 |  |  |
| miR-576-5p | -1.86 | miR-215 | -1.80 | miR-651 | -3.12 |  |  |
| miR-193a-5p | -1.89 | miR-425-5p | -1.85 | miR-375 | -3.15 |  |  |
| miR-215 | -1.91 | miR-625-3p | -1.89 | miR-135a-5p | -3.27 |  |  |
| miR-124-3p | -1.93 | miR-205-5p | -1.89 | miR-708-5p | -3.51 |  |  |
| miR-574-3p | -1.98 | miR-26a-5p | -1.90 | miR-421 | -3.67 |  |  |
| miR-302c-5p | -2.01 | miR-582-5p | -1.97 | miR-34b-3p | -3.79 |  |  |
| miR-142-3p | -2.11 | miR-197-3p | -2.07 | miR-219-5p | -3.82 |  |  |
| miR-361-5p | -2.16 | miR-26b-5p | -2.11 | miR-550a-5p | -4.12 |  |  |
| miR-486-5p | -2.23 | miR-219-5p | -2.13 | miR-203a | -4.37 |  |  |
| miR-185-3p | -2.26 | miR-224-5p | -2.18 | miR-330-3p | -4.61 |  |  |
| miR-196a-5p | -2.35 | miR-181a-3p | -2.31 | miR-200c-3p | -6.61 |  |  |
| miR-140-3p | -2.35 | let-7d-3p | -2.39 | miR-141-3p | -6.69 |  |  |
| miR-100-5p | -2.40 | miR-452-5p | -2.47 | miR-205-5p | -10.23 |  |  |
| miR-326 | -2.45 | miR-100-5p | -2.59 |  |  |  |  |
| miR-452-5p | -2.47 | miR-361-3p | -2.60 |  |  |  |  |
| miR-425-3p | -2.57 | miR-95 | -2.74 |  |  |  |  |
| miR-548b-3p | -2.65 | miR-99a-5p | -2.79 |  |  |  |  |

|  |  |  |  |
| --- | --- | --- | --- |
| miR-95 | -2.74 | miR-455-5p | -2.86 |
| miR-29b-2-5p | -2.75 | miR-181b-5p | -3.00 |
| miR-654-5p | -2.82 | miR-570-3p | -3.01 |
| miR-21-3p | -2.90 | miR-186-5p | -3.04 |
| miR-324-3p | -2.90 | miR-21-3p | -3.16 |
| miR-421 | -2.99 | miR-425-3p | -3.19 |
| miR-615-3p | -3.02 | miR-140-3p | -3.87 |
| miR-33b-5p | -3.08 | miR-18a-3p | -3.94 |
| miR-651 | -3.12 | miR-503-5p | -3.95 |
| miR-26a-5p | -3.19 | miR-421 | -3.96 |
| miR-589-5p | -3.25 | miR-576-3p | -4.01 |
| miR-373-5p | -3.26 | miR-545-3p | -4.54 |
| let-7d-3p | -3.33 | miR-146b-5p | -6.19 |
| miR-223-3p | -3.39 | miR-663a | -7.62 |
| miR-30c-2-3p | -3.47 |  |  |
| miR-18a-3p | -3.67 |  |  |
| miR-34b-3p | -3.79 |  |  |
| miR-625-3p | -3.80 |  |  |
| miR-181a-3p | -3.94 |  |  |
| miR-374b-5p | -4.09 |  |  |
| miR-150-5p | -4.71 |  |  |
| miR-186-5p | -5.79 |  |  |
| miR-874 | -7.04 |  |  |
| miR-146b-5p | -8.33 |  |  |
| miR-663a | -9.69 |  |  |

**AE:** AURKB MCF10A EVs, **ME:** MYC MCF10A EVs, **HE:** HRAS MCF10A EVs, **CE:** CCDN1 MCF10A EVs, **PE:** Puro/Blast (control) MCF10A EVs

**Supplementary Table 5.**
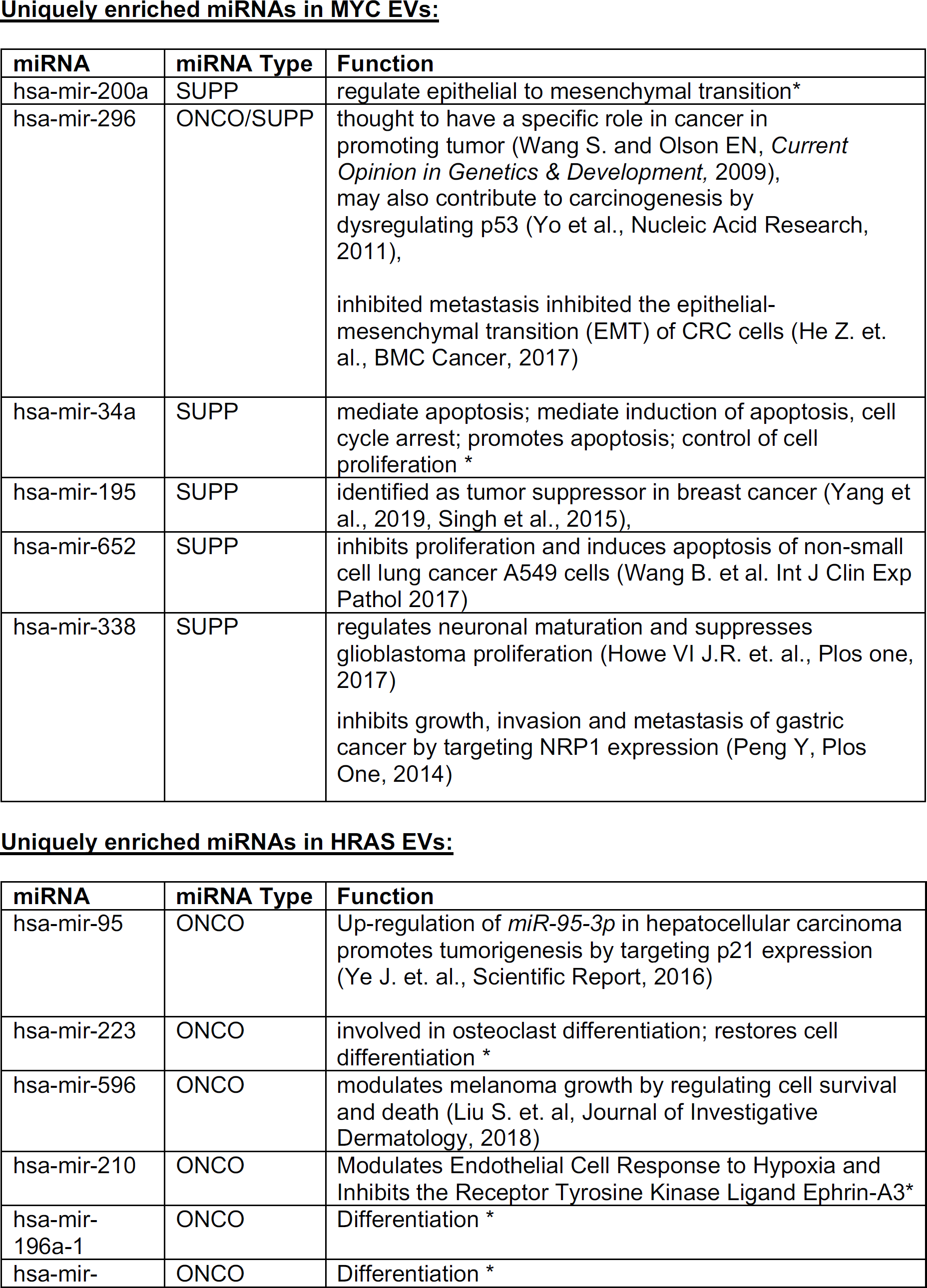
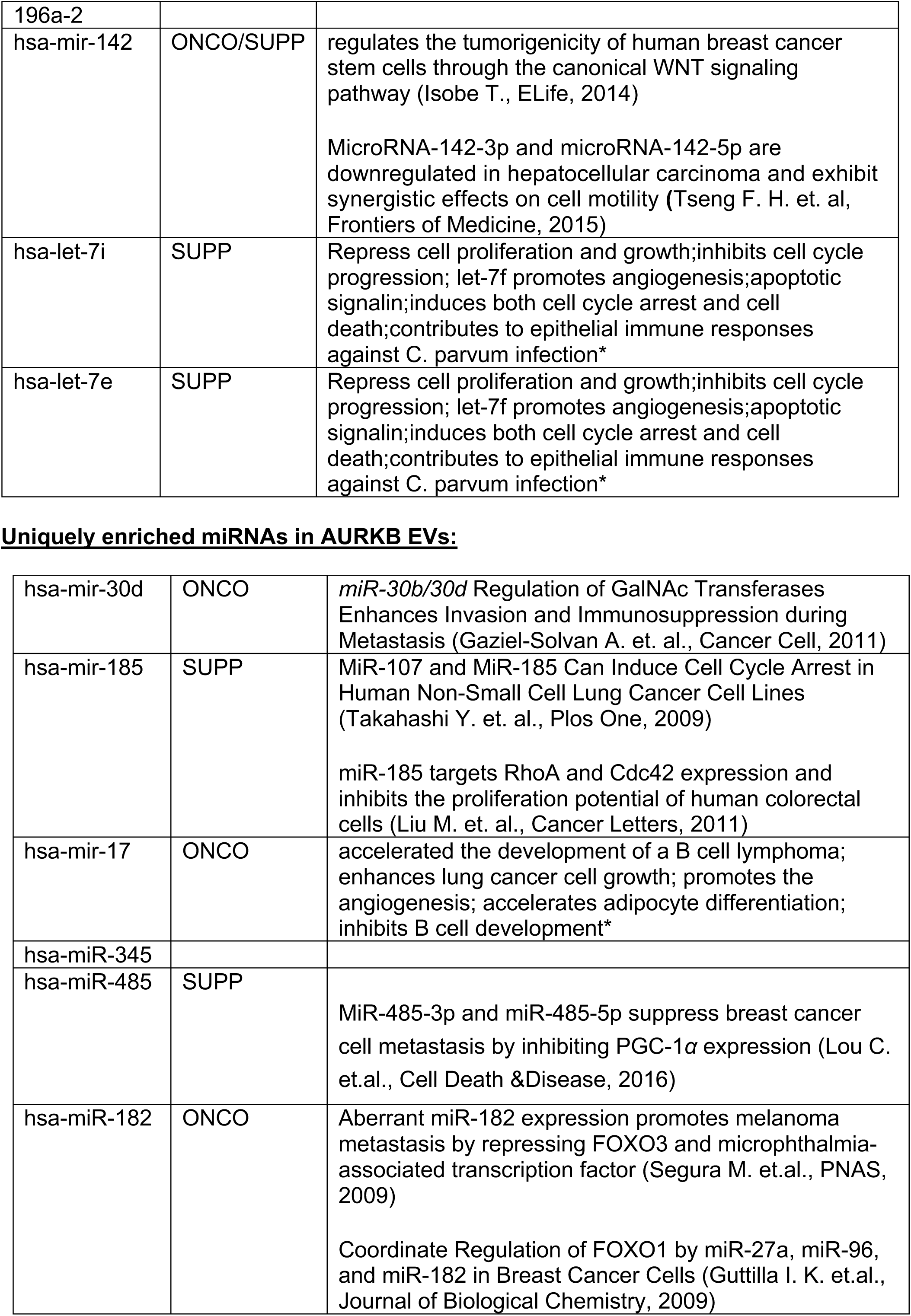
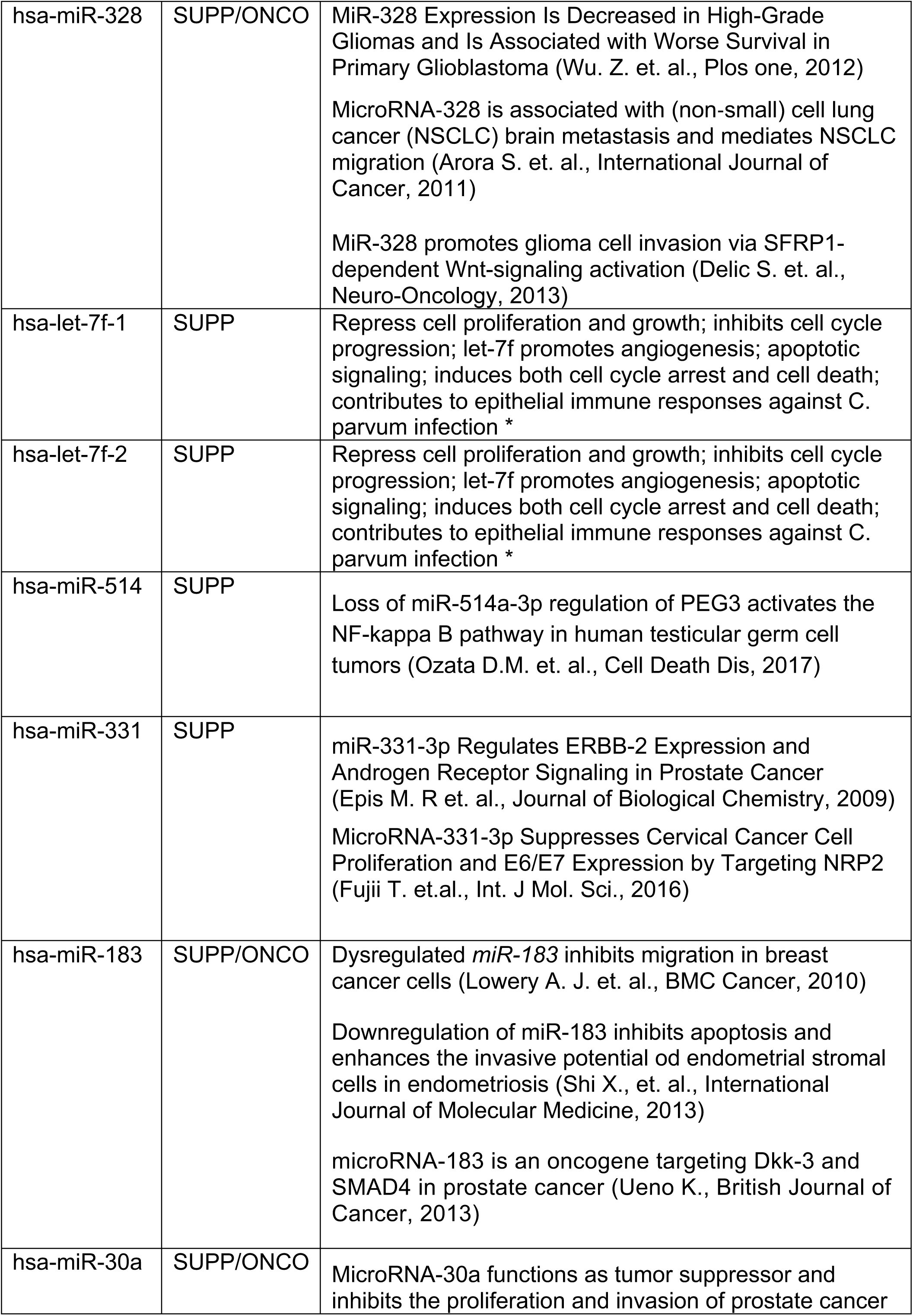
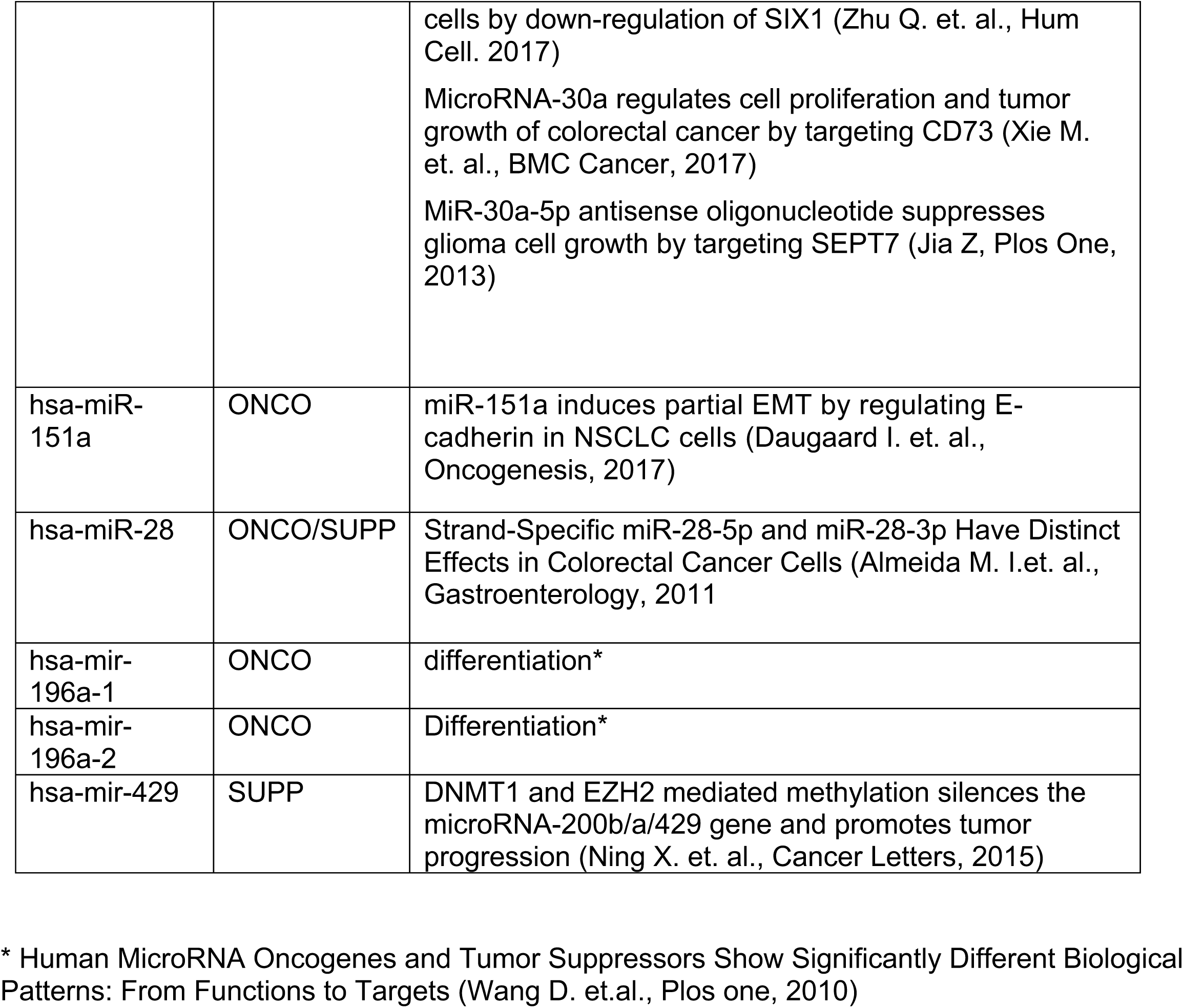
Functions of miRNAs identified uniquely in each oncogene.

